# Identification and characterization of *ethR* and *ethA* genes impacting the sensitivity of *Mycobacterium abscessus* to ethionamide

**DOI:** 10.64898/2026.02.08.704650

**Authors:** Xiaofan Zhang, Lijie Li, Ziwen Lu, Zhaohui Duan, H.M. Adnan Hameed, Belachew Aweke Mulu, Cuiting Fang, Xirong Tian, Xinyue Wang, Hongyi Chen, Liqiang Feng, Matthew B. McNeil, Xindong Liu, Shuai Wang, Tianyu Zhang

**Affiliations:** State Key Laboratory of Respiratory Disease, Guangzhou Institutes of Biomedicine and Health, Chinese Academy of Sciences, Guangzhou 510530, China; University of Chinese Academy of Sciences, Beijing 100049, China; Department of Clinical Laboratory, Sun Yat-sen Memorial Hospital of Sun Yat-sen University, Guangzhou 510120, China; School of Life Sciences, University of Science and Technology of China, Hefei 230027, China; China-New Zealand Joint Laboratory on Biomedicine and Health, Guangzhou Institutes of Biomedicine and Health, Chinese Academy of Sciences, Guangzhou 510530, China; Guangdong-Hong Kong-Macao Joint Laboratory of Respiratory Infectious Diseases, Guangzhou Institutes of Biomedicine and Health, Chinese Academy of Sciences, Guangzhou 510530, China; Guangzhou Medical University-Guangzhou Institutes of Biomedicine and Health Joint School of Life Sciences, Guangzhou Medical University, Guangzhou 511436, China; State Key Laboratory of Respiratory Disease, Guangzhou Chest Hospital, Guangzhou 510095, China; Guangzhou National Laboratory, Guangzhou 510005, China; West China School of Public Health, Sichuan University, Chengdu 610041, China; Department of Microbiology and Immunology, School of Biomedical Sciences, University of Otago, Dunedin 9054, New Zealand; Maurice Wilkins Centre for Molecular Biodiscovery, University of Auckland, Auckland 1042, New Zealand; Chinese Center for Disease Control and Prevention, National Pathogen Resource Center, Beijing 102206, China

**Keywords:** *Mycobacterium abscessus*, ethionamide, intrinsic resistance, *ethR*, *ethA*

## Abstract

*Mycobacterium abscessus* is a rapidly growing non-tuberculous mycobacterium with rising global incidence. This pathogen exhibits intrinsic resistance to most antibiotics, presenting a major public health threat. Ethionamide (ETH) requires bioactivation by monooxygenase EthA to form the active ETH-NAD adduct. We previously identified MAB_3513 (NudC) as a phosphohydrolase that confers intrinsic resistance to *M. abscessus* by hydrolyzing this adduct. However, deletion of *nudC* results in only partial susceptibility to ETH, indicating the existence of additional resistance mechanisms. This study identified MAB_0984 as the EthR regulator in *M. abscessus*. Deletion of *ethR* in a *nudC* knockout background (ΔΔethR) significantly enhanced ETH-NAD adduct accumulation, leading to hypersusceptibility to ETH. Notably, the ΔethR mutant exhibited higher susceptibility than ΔnudC, demonstrating that EthR is a more dominant mediator of ETH resistance than NudC. Furthermore, MAB_0985 (EthA1) and MAB_0103 (EthA2) were identified as the primary EthAs in *M. abscessus*. Deletion of either gene alone or in combination in the ΔnudC reduced adduct formation and increased resistance, while the triple mutant ΔΔethA1ΔethA2 restored wild-type resistance. Using an intergenic region-eGFP reporter system and quantitative reverse transcription-PCR, we demonstrated that EthR confers resistance by specifically suppressing *ethA1* expression in *M. abscessus*. The *Mycobacterium tuberculosis* EthR inhibitor BDM31343 could boost the efficacy of ETH against *M. abscessus* by inhibiting EthR. Collectively, this study identified the *ethR* and *ethA* genes in *M. abscessus* for the first time and elucidated their role in mediating resistance to ETH. Therefore, EthR is a promising target for potentiating the efficacy of ETH against *M. abscessus*.

**Impact statement:** *Mycobacterium abscessus* constitutes an escalating global health threat, primarily due to its intrinsic resistance to most antibiotics. This study identifies MAB_0984 (EthR) as a dominant resistance determinant that exerts a more profound impact on ETH susceptibility than the previously characterized NudC. We demonstrate that EthR mediates resistance by specifically repressing the expression of EthA1 (MAB_0985), one of the primary monooxygenases responsible for ETH bioactivation. The EthR^Mtb^ inhibitor BDM31343 potentiated the activity of ETH against *M. abscessus* through inhibition of the EthR. These findings elucidated the mechanism of ETH resistance in *M. abscessus*, identifying EthR as a promising target for boosting the efficacy of ETH against *M. abscessus*.

## INTRODUCTION

*Mycobacterium abscessus* (*M. abscessus*) is a rapidly growing nontuberculous mycobacterium (NTM) that causes severe pulmonary, cutaneous, and subcutaneous infections in humans. Over the past two decades, the global incidence of NTM infections has increased markedly across diverse geographic regions, including North America, Europe, Asia, and Australia^1^. In developed countries, the incidence and prevalence of NTM pulmonary disease now exceed those of tuberculosis^2^. Among NTMs, *M. abscessus* is the second most common pathogen causing NTM disease and the most prevalent rapidly growing species, with disease burden particularly pronounced in an aging populations^1^. As an environmental opportunistic pathogen, *M. abscessus* is widely distributed in the environment, and recent studies have also confirmed its potential for person-to-person transmission^3^. Together, these factors underscore *M. abscessus* as an emerging public health threat that demands urgent attention.

The clinical management of *M. abscessus* infections is severely compromised by its extensive intrinsic and acquired resistance to antibiotics. This multidrug resistance is driven by multiple mechanisms, including a lipid-rich impermeable cell wall, active drug efflux pumps, antibiotic-modifying enzymes, and polymorphisms in drug target genes^4^. Consequently, the extensive nature of drug resistance, combined with the lack of consistently effective treatment regimens, makes it extremely difficult to treat these infections. Treatment outcomes for *M. abscessus* infections are often poor, with combination antibiotic therapy achieving success in only 30-50% of patients, and surgical resection is frequently required as an adjunctive intervention to control the infection^5^. Current guidelines recommend prolonged treatment regimens for *M. abscessus* pulmonary disease, typically involving 6-12 months of combined oral and intravenous antibiotics^6^, followed by continuation of therapy for at least 12 months after sputum culture conversion^7^. Oral regimens generally comprises one or two drugs, such as a macrolide, linezolid, or clofazimine with fluoroquinolones are occasionally incorporated. Intravenous therapy commonly involves two or more drugs including amikacin (AMK), tigecycline, imipenem, or cefoxitin. Although newer agents such as bedaquiline (BDQ) and its next-generation derivatives TBAJ-587 and TBAJ-876 have demonstrated promising *in vitro* and *in vivo* activity against *M. abscessus*^8,9^, treatment success remains suboptimal, particularly in chronic, refractory pulmonary disease, which is frequently associated with relapse and increased mortality. These challenges highlight an urgent need to elucidate the molecular basis of antibiotic resistance in *M. abscessus* and to identify novel therapeutic targets.

Ethionamide (ETH) is a second-line antitubercular drug widely used for the treatment of multidrug-resistant tuberculosis. As a prodrug and structural analog of isoniazid (INH), ETH is activated by the bacterial monooxygenases to form an active ETH-NAD adduct. This adduct inhibits the enoyl-acyl carrier protein reductase (InhA) in the fatty acid synthase II (FAS-II) pathway, disrupting mycolic acid synthesis and ultimately causing mycobacterial cell death. Three monooxygenases identified in *Mycobacterium tuberculosis*, EthA^Mtb^ (Rv3854)^10^, EthA2^Mtb^ (Rv0077)^11^, and MymA (Rv3083)^12^, have been shown to activate ETH. The expression of these enzymes is tightly regulated by the transcriptional regulators EthR^Mtb^ (Rv3855)^10^, EthR2^Mtb^ (Rv0078)^11^, and VirS (Rv3082)^12^, respectively. EthR^Mtb^ and EthR2^Mtb^ are members of the TetR family of repressors, whereas VirS belongs to the AraC/XylS family of transcriptional factors. Overexpression of *ethA*^Mtb^, *ethA2*^Mtb^, or *mymA* increased *M. tuberculosis* susceptibility to ETH^10–12^. Furthermore, small-molecule inhibitors targeting these repressors, BDM41906, SMARt420, and SMARt751, have been shown to derepress monooxygenase expression, thereby markedly potentiating ETH activity both *in vitro* and *in vivo*^11,13,14^. In contrast to *M. tuberculosis*, *M. abscessus* exhibits high-level intrinsic resistance to ETH, and the precise molecular mechanism underlying this resistance remains incompletely characterized. Our prior investigation demonstrated that the *MAB_3513c* gene in *M. abscessus* encodes NudC, a pyrophosphohydrolase enzyme that confers ETH resistance by hydrolyzing the ETH-NAD adduct^15^. Although the *nudC* knockout reduced the minimal inhibitory concentration (MIC) of ETH against *M. abscessus* from 256 μg/mL to 16 μg/mL, the value remains much higher than the MIC to *M. tuberculosis*, indicating additional resistance mechanisms in *M. abscessus*.

This study is the first to identify MAB_0984 as the functional EthR homolog in *M. abscessus*, which regulates EthA1 (MAB_0985) expression and thereby suppresses ETH-NAD adduct formation. Our findings reveal that impaired EthR-regulated activation is a more critical determinant of intrinsic ETH resistance in *M. abscessus* than the previously identified NudC-mediated detoxification pathway. Additionally, our study demonstrates that MAB_0985 and MAB_0103 function as the principal EthAs in *M. abscessus*, mediating ETH activation and the subsequent formation of ETH-NAD adduct. The small-molecule inhibitor BDM31343, targeting EthR in *M. tuberculosis*, enhanced the efficacy of ETH against *M. abscessus*. Collectively, our findings elucidate the mechanism of ETH resistance in *M. abscessus*, thereby identifying EthR as a promising therapeutic target.

## RESULTS

### Deletion of *MAB_0984c* encoding a homolog of the EthR^Mtb^ increased *M. abscessus* susceptibility to ETH

MAB_0984 was identified as the EthR homolog in *M. abscessus*, exhibiting 50.23% amino acid sequence identity with EthR^Mtb^ (Figure S1). Both proteins belong to the TetR transcriptional regulator family. In contrast, no homologs exhibiting > 40% amino acid sequence identity to EthR2^Mtb^ or VirS were detected in *M. abscessus*, suggesting that MAB_0984 represents the primary EthR-like regulator in *M. abscessus*.

To investigate its role in ETH resistance, CRISPR-assisted non-homologous end-joining (NHEJ) technology^16^ was used to knock out *MAB_0984c* (*ethR*) in ΔnudC. A frameshift mutation was generated near the 5’ end of the *ethR* by introducing a 4-bp deletion (nucleotides 191-194), which was confirmed by Sanger sequencing (Figure S2A,B). The resulting double mutant, ΔnudCΔethR (hereafter referred to as ΔΔethR), exhibited extreme sensitivity to ETH, with a MIC of 0.125 μg/mL (Table 1). This represents a 128-fold reduction relative to the MIC to ΔnudC (16 μg/mL) and a 2048-fold reduction relative to the MIC to the WT strain (256 μg/mL). In contrast, their MICs to rifabutin (RFB), moxifloxacin (MXF), INH, AMK, and BDQ remained unchanged. This finding suggests that EthR constitutes an additional critical factor, beyond NudC, conferring ETH resistance in *M. abscessus.* To further investigate this, we engineered an *ethR* knockout (ΔethR) strain in WT through an in-frame deletion of 240 bp beginning at nucleotide position 46 of the coding sequence, which was verified by Sanger sequencing (Figure S2C,D). The MIC of ETH to ΔethR was 4 μg/mL, representing a 64-fold reduction compared to the WT and a 4-fold reduction relative to the ΔnudC (Table 1). The ETH resistance was successfully restored in both the ΔΔethR and ΔethR upon complementation with *ethR* or *ethR*^Mtb^ (Table 1), confirming their functional homology. On agar plates, ΔΔethR and ΔethR exhibited sensitivity to ETH at concentrations of 0.25 μg/mL and 4 μg/mL (Figure 1A,B), respectively, in agreement with the MIC values. Furthermore, upon exposure to 256 μg/mL ETH, the accumulation of the ETH-NAD adduct in the ΔΔethR was significantly increased, reaching a level 2.5-fold greater than that in the ΔnudC (Figure 1C). Knockout of *ethR* in both ΔnudC and WT did not alter their respective growth rates (Figure S3). These results indicate that MAB_0984 is the functional EthR in *M. abscessus*, contributing to intrinsic ETH resistance alongside NudC. Notably, EthR-mediated resistance is more pronounced than that attributed to NudC in *M. abscessus*.

**Table 1.** The susceptibility of *M. abscessus ethR* knockout strains in a *nudC* deletion and wild-type background, as well as their complementary strains, to ETH and INH.

| *M. abscessus* strains | Antibiotics/ MICs<sup>a</sup> (μg/mL) |  |  |  |  |  |
| --- | --- | --- | --- | --- | --- | --- |
|  | ETH | INH | AMK | RFB | BDQ | MXF |
| Δ*nudC* | 16 | 1 | 16 | 8 | 1 | 16 |
| ΔΔ*ethR* | 0.125 | 1 | 16 | 8 | 1 | 16 |
| ΔΔ*ethR*<sup>CMab</sup> | 16 | 1 | / | / | / | / |
| ΔΔ*ethR*<sup>CMtb</sup> | 32 | 1 | / | / | / | / |
| ΔΔ*ethR*<sup>CEV</sup> | 0.125 | 1 | / | / | / | / |
| WT | 256 | > 512 | 16 | 8 | 1 | 16 |
| Δ*ethR* | 4 | > 512 | 16 | 8 | 1 | 16 |
| Δ*ethR*<sup>CMab</sup> | 256 | > 512 | / | / | / | / |
| Δ*ethR*<sup>CMtb</sup> | 256 | > 512 | / | / | / | / |
| Δ*ethR*<sup>CEV</sup> | 4 | > 512 | / | / | / | / |

<sup>a</sup> Broth microdilution method was used to determine the MICs. Δ*nudC*: *M. abscessus nudC* knockout strain; ΔΔ*ethR*: *ethR* and *nudC* double knockout strain; ΔΔ*ethR*<sup>CMab</sup>, ΔΔ*ethR*<sup>CMtb</sup>, ΔΔ*ethR*<sup>CEV</sup>: ΔΔ*ethR* complemented with *ethR*<sup>Mab</sup>, *ethR*<sup>Mtb</sup>, or with the empty vector, respectively. WT: wild-type *M. abscessus*; Δ*ethR*: *ethR* knockout strain; Δ*ethR*<sup>CMab</sup>, Δ*ethR*<sup>CMtb</sup>, Δ*ethR*<sup>CEV</sup>: Δ*ethR* complemented with *ethR*<sup>Mab</sup>, *ethR*<sup>Mtb</sup>, or with the empty vector, respectively.

**Figure 1.**
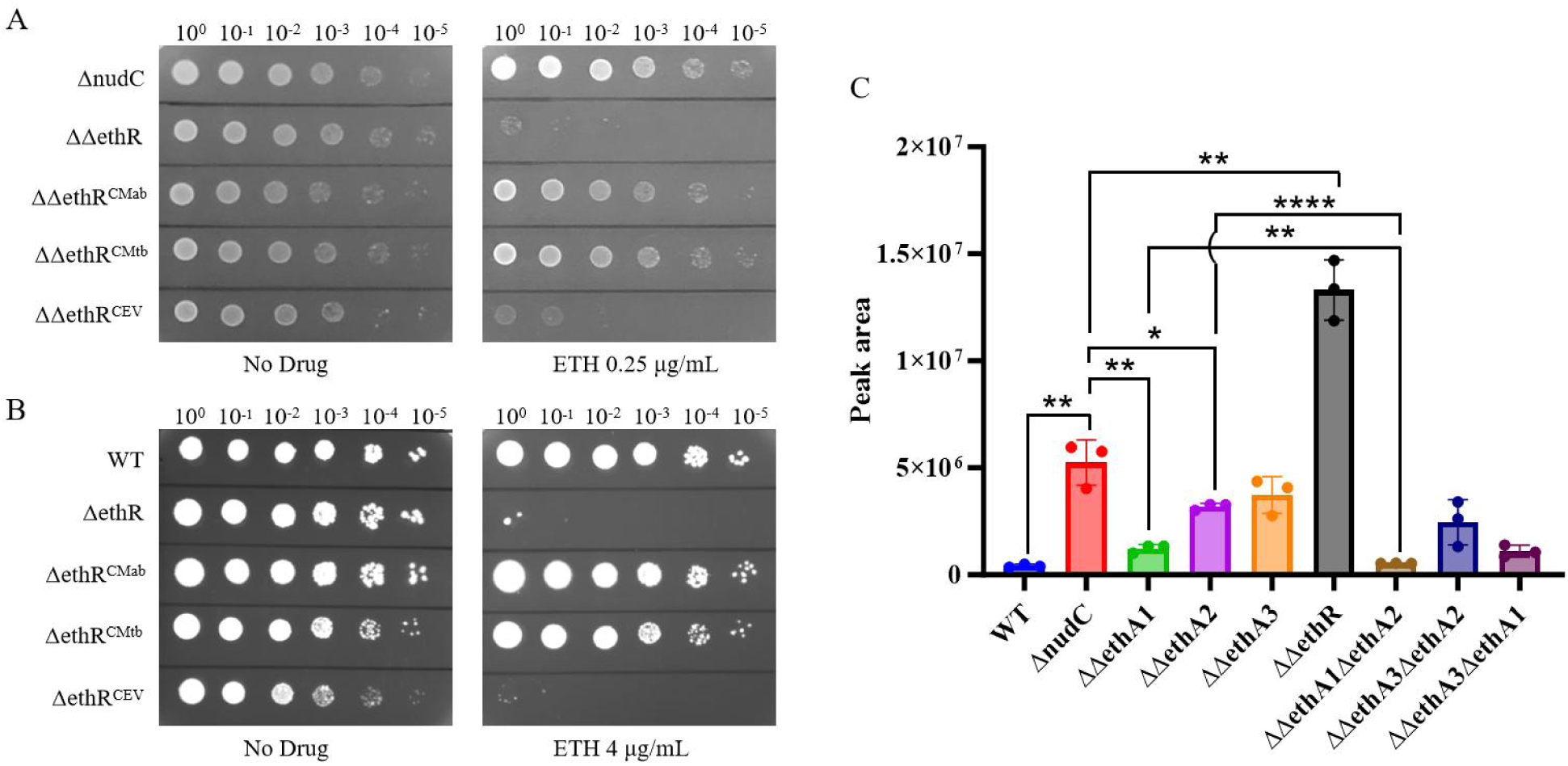
Knockout of *ethR* (*MAB_0984c*) significantly enhances the susceptibility to ETH by promoting the accumulation of the ETH-NAD adduct within *M. abscessus*. (A) Drug susceptibility testing of the *ethR* and *nudC* double knockout strain to ETH. ΔnudC: *M. abscessus nudC* knockout strain; ΔΔethR: *ethR* and *nudC* double knockout strain; ΔΔethR^CMab^, ΔΔethR^CMtb^, ΔΔethR^CEV^: ΔΔethR complemented with *ethR*^Mab^, *ethR*^Mtb^, or with the empty vector, respectively. (B) Drug susceptibility testing of the *ethR* knockout strain to ETH. WT: wild-type *M. abscessus*; ΔethR: *ethR* knockout strain; ΔethR^CMab^, ΔethR^CMtb^, ΔethR^CEV^: ΔethR complemented with *ethR*^Mab^, *ethR*^Mtb^, or with the empty vector, respectively. (C) Quantification of the ETH-NAD adduct in the various *M. abscessus* strains. ΔΔethA1: *ethA1* (*MAB_0985*) and *nudC* double knockout strain; ΔΔethA2: *ethA2* (*MAB_0103*) and *nudC* double knockout strain; ΔΔethA3: *ethA3* (*MAB_3967*) and *nudC* double knockout strain; ΔΔethA1ΔethA2: *ethA1*, *ethA2*, and *nudC* triple knockout strain; ΔΔethA3ΔethA2: *ethA3*, *ethA2*, and *nudC* triple knockout strain; ΔΔethA3ΔethA1: *ethA3*, *ethA1* and *nudC* triple knockout strain.

### Deletion of *MAB_0985* and *MAB_0103* encoding homologs of the EthA^Mtb^ enhanced *M. abscessus* resistance to ETH

Given that MAB_0984 functions as EthR in *M. abscessus* and that EthR acts by regulating EthA expression, a critical question arises: which gene serves as the functional EthA target in *M. abscessus*? BLASTp analysis identified MAB_0985, MAB_0103, and MAB_3967 in *M. abscessus* as potential EthA homologs, exhibiting 63.46% amino acid sequence identity to the known EthA^Mtb^ (Figure S4). Considering the intrinsic resistance of WT to ETH, we initially employed CRISPR-NHEJ to knockout three potential *ethA* genes in the ΔnudC (Figure S5). The double knockout strains ΔnudCΔ0985 (ΔΔ0985) and ΔnudCΔ0103 (ΔΔ0103) demonstrated modestly elevated ETH resistance, with MICs increasing 4-fold and 2-fold, respectively (Table 2). In contrast, their MICs for INH, AMK, RFB, BDQ, and MXF remained unchanged. ETH susceptibility was restored upon individual reintroduction of *MAB_0985*, *MAB_0103*, or *ethA*^Mtb^ into their respective mutants (Table S1). The enhanced ETH resistance in mutant strains was directly observable on solid plates, with ΔΔ0985 exhibiting growth at 32 μg/mL and ΔΔ0103 at 16 μg/mL, indicating modestly increased resistance (Figure 2). The ΔΔ3967 mutant, diluted 10,000-fold on plates containing 8 μg/mL of ETH, exhibited slightly higher resistance than ΔnudC, though this difference was not statistically significant.(Figure S6).

**Table 2.**
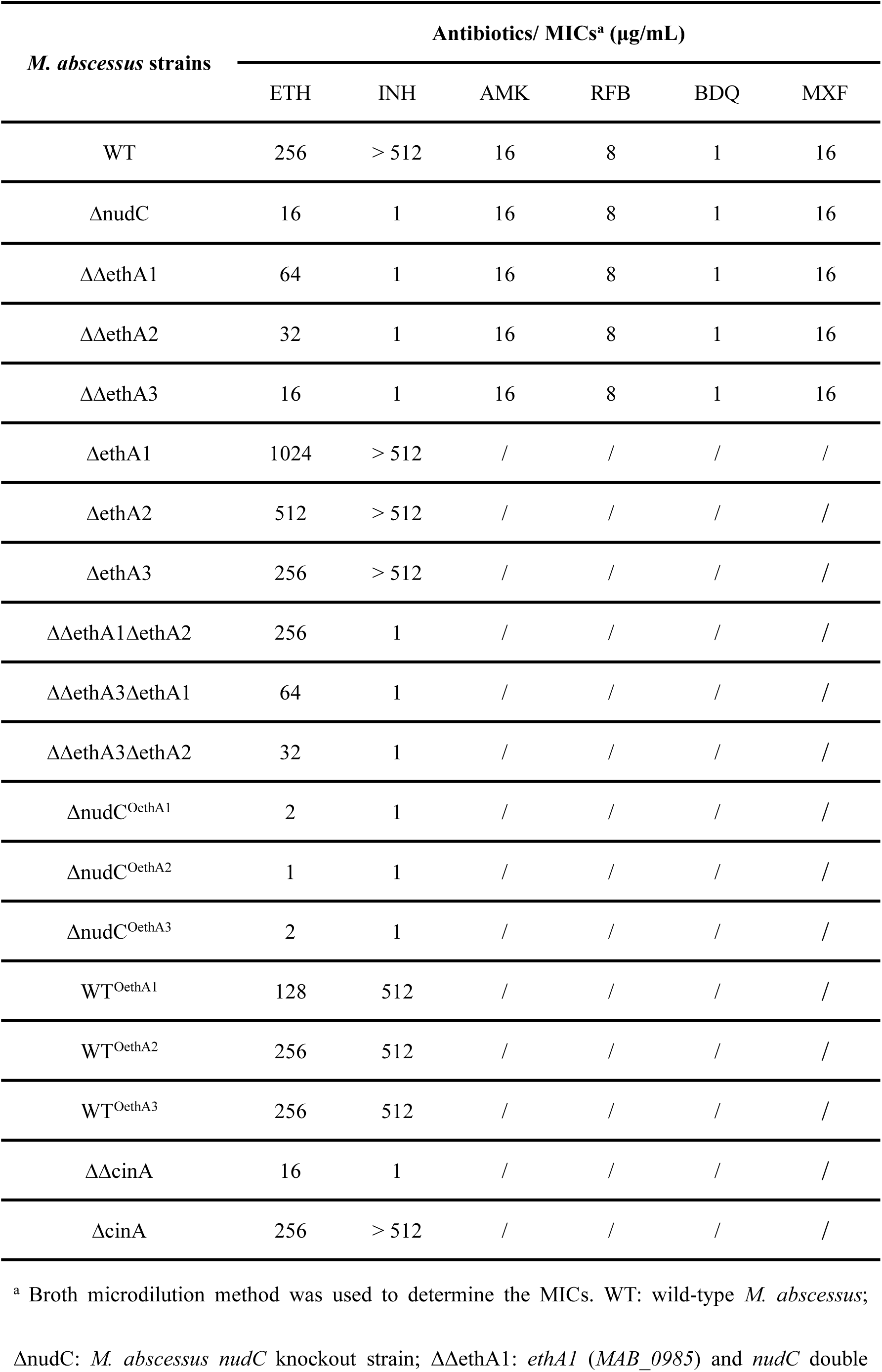
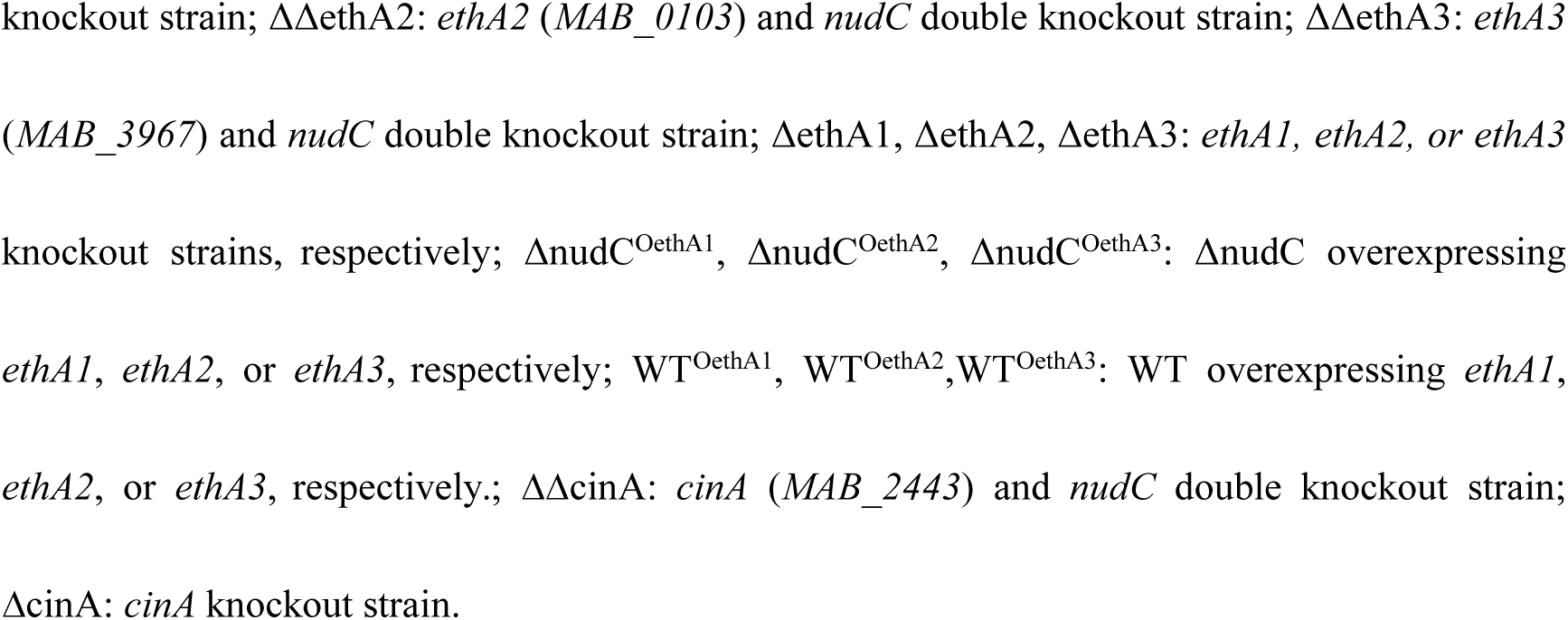
The susceptibility of different *M. abscessus* strains to various drugs

| *M. abscessus* strains | Antibiotics/ MICs<sup>a</sup> (µg/mL) |  |  |  |  |  |
| --- | --- | --- | --- | --- | --- | --- |
|  | ETH | INH | AMK | RFB | BDQ | MXF |
| WT | 256 | > 512 | 16 | 8 | 1 | 16 |
| ΔnudC | 16 | 1 | 16 | 8 | 1 | 16 |
| ΔΔethA1 | 64 | 1 | 16 | 8 | 1 | 16 |
| ΔΔethA2 | 32 | 1 | 16 | 8 | 1 | 16 |
| ΔΔethA3 | 16 | 1 | 16 | 8 | 1 | 16 |
| ΔethA1 | 1024 | > 512 | / | / | / | / |
| ΔethA2 | 512 | > 512 | / | / | / | / |
| ΔethA3 | 256 | > 512 | / | / | / | / |
| ΔΔethA1ΔethA2 | 256 | 1 | / | / | / | / |
| ΔΔethA3ΔethA1 | 64 | 1 | / | / | / | / |
| ΔΔethA3ΔethA2 | 32 | 1 | / | / | / | / |
| ΔnudC<sup>OethA1</sup> | 2 | 1 | / | / | / | / |
| ΔnudC<sup>OethA2</sup> | 1 | 1 | / | / | / | / |
| ΔnudC<sup>OethA3</sup> | 2 | 1 | / | / | / | / |
| WT<sup>OethA1</sup> | 128 | 512 | / | / | / | / |
| WT<sup>OethA2</sup> | 256 | 512 | / | / | / | / |
| WT<sup>OethA3</sup> | 256 | 512 | / | / | / | / |
| ΔΔcinA | 16 | 1 | / | / | / | / |
| ΔcinA | 256 | > 512 | / | / | / | / |

<sup>a</sup> Broth microdilution method was used to determine the MICs. WT: wild-type *M. abscessus*;
ΔnudC: *M. abscessus nudC* knockout strain; ΔΔethA1: *ethA1* (*MAB\_0985*) and *nudC* double
knockout strain; $\Delta\Delta\text{ethA2}$ : *ethA2* (*MAB\_0103*) and *nudC* double knockout strain; $\Delta\Delta\text{ethA3}$ : *ethA3* (*MAB\_3967*) and *nudC* double knockout strain; $\Delta\text{ethA1}$ , $\Delta\text{ethA2}$ , $\Delta\text{ethA3}$ : *ethA1*, *ethA2*, or *ethA3* knockout strains, respectively; $\Delta\text{nudC}^{\text{OethA1}}$ , $\Delta\text{nudC}^{\text{OethA2}}$ , $\Delta\text{nudC}^{\text{OethA3}}$ : $\Delta\text{nudC}$ overexpressing *ethA1*, *ethA2*, or *ethA3*, respectively; $\text{WT}^{\text{OethA1}}$ , $\text{WT}^{\text{OethA2}}$ , $\text{WT}^{\text{OethA3}}$ : WT overexpressing *ethA1*, *ethA2*, or *ethA3*, respectively.; $\Delta\Delta\text{cinA}$ : *cinA* (*MAB\_2443*) and *nudC* double knockout strain; $\Delta\text{cinA}$ : *cinA* knockout strain.

**Figure 2.**
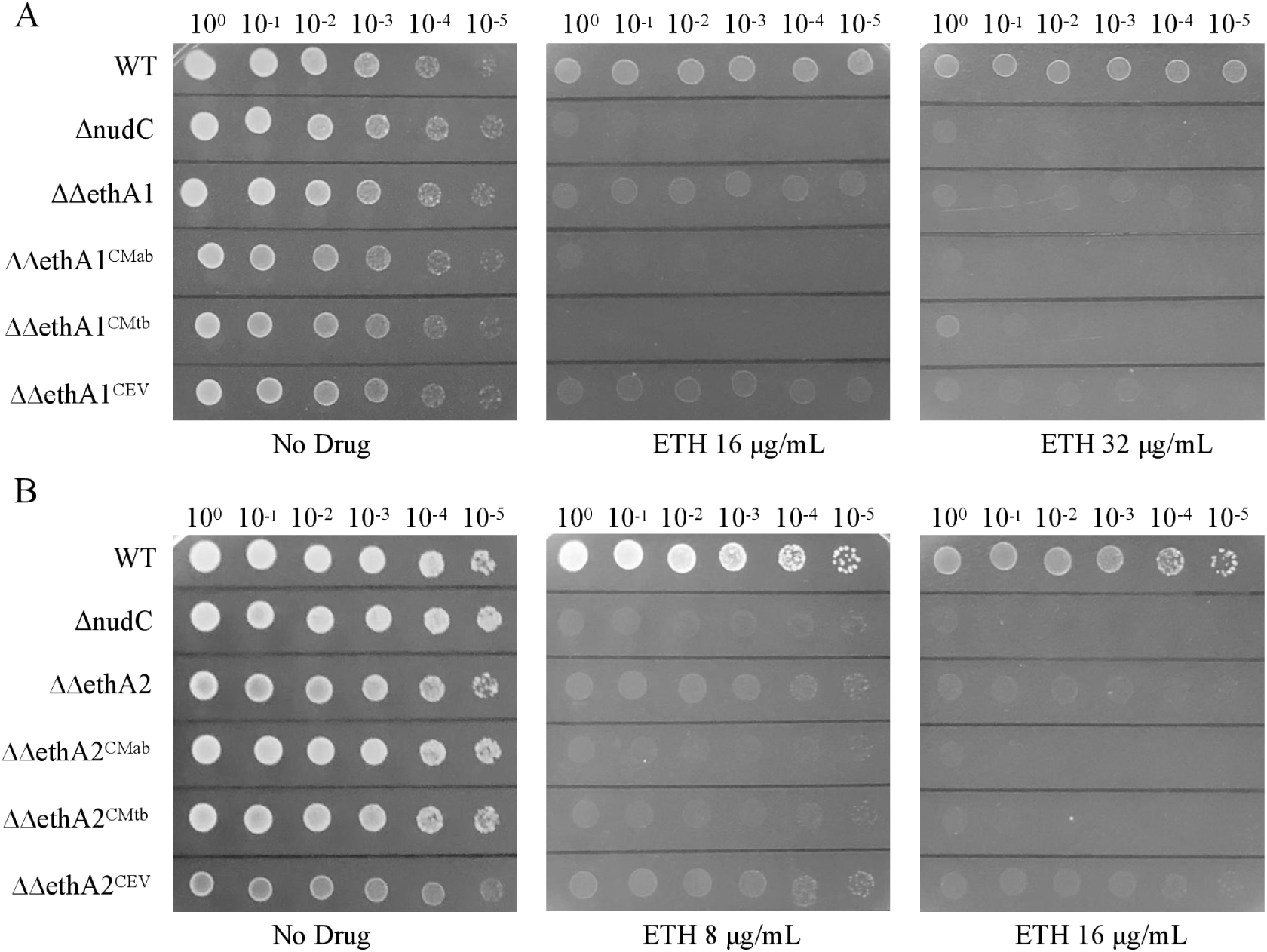
Knockout of *ethA1* (*MAB_0985*) or *ethA2* (*MAB_0103*) in the *nudC* knockout strain increases resistance to ETH. (A) Drug susceptibility testing of the *ethA1* and *nudC* double knockout strain to ETH. WT: wild-type *M. abscessus*; ΔnudC: *M. abscessus nudC* knockout strain; ΔΔethA1: *ethA1* and *nudC* double knockout strain; ΔΔethA1^CMab^, ΔΔethA1^CMtb^, ΔΔethA1^CEV^: ΔΔethA1 complemented with *ethA1*^Mab^, *ethA*^Mtb^, or with the empty vector, respectively. (B) Drug susceptibility testing of the *ethA2* and *nudC* double knockout strain to ETH. ΔΔethA2: *ethA2* and *nudC* double knockout strain; ΔΔethA2^CMab^, ΔΔethA2^CMtb^, ΔΔethA2^CEV^: ΔΔethA2 complemented with *ethA2*^Mab^, *ethA*^Mtb^, or with the empty vector, respectively.

To further investigate the functions of *MAB_0985*, *MAB_0103*, and *MAB_3967*, we generated single knockout mutants of each gene in the WT background (Figure S7). Drug susceptibility testing (DST) demonstrated that Δ0985 and Δ0103 exhibited increased resistance to ETH compared to the WT (Table 2). The MICs to ETH rose 4-fold in the Δ0985 and 2-fold in the Δ0103, whereas the MIC of INH remained unchanged (Table 2). INH is specifically activated by the catalase-peroxidase KatG to form the INH-NAD adduct, which also subsequently inhibits the InhA. Consequently, deletion of ethAs did not affect susceptibility to INH in the ΔnudC. Enhanced ETH resistance was directly demonstrated on solid medium, as evidenced by the growth of the Δ0985 growing at 512 μg/mL and Δ0103 at 256 μg/mL of ETH, consistent with a resistant phenotype (Figure 3A). The results demonstrate that deletion of *MAB_0985* or *MAB_0103* increased *M. abscessus* resistance to ETH.

**Figure 3.**
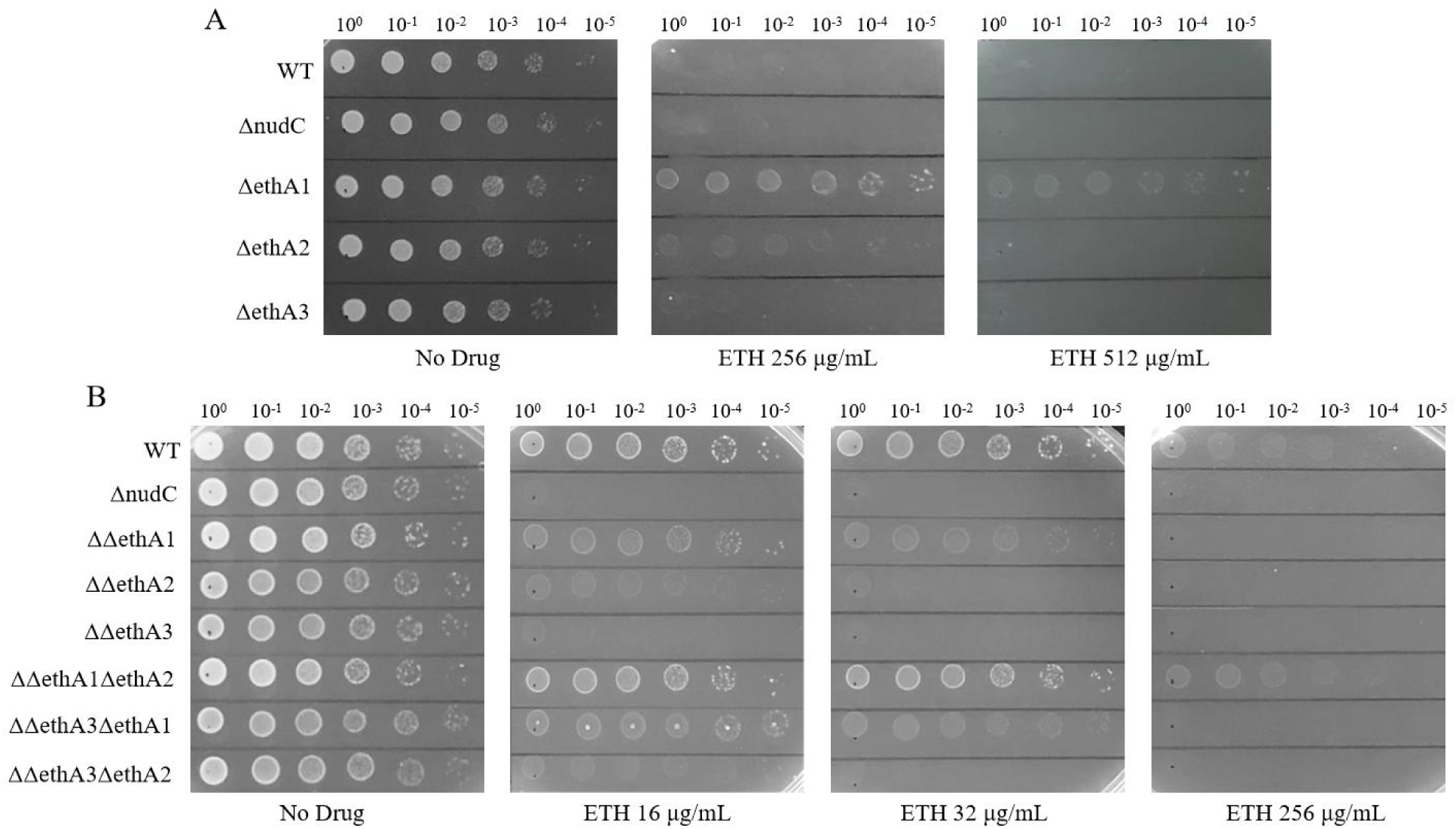
Knockout of *ethA1* (*MAB_0985*) or *ethA2* (*MAB_0103*) in wild-type *M. abscessus* increases resistance to ETH, and simultaneous knockout of *ethA1* and *ethA2* in the *nudC* knockout strain restores wild-type resistance levels. (A) Drug susceptibility testing of *ethA1*, *ethA2*, and *ethA3* (*MAB_3967*) knockout strain to ETH. WT: wild-type *M. abscessus*; ΔnudC: *M. abscessus nudC* knockout strain; ΔethA1: *ethA1* knockout strain; ΔethA2: *ethA2* knockout strain; ΔethA3: *ethA3* knockout strain. (B) Drug susceptibility testing of the triple knockout strain to ETH. ΔΔethA1: *ethA1* and *nudC* double knockout strain; ΔΔethA2: *ethA2* and *nudC* double knockout strain; ΔΔethA3: *ethA3* and *nudC* double knockout strain; ΔΔethA1ΔethA2: *ethA1*, *ethA2*, and *nudC* triple knockout strain; ΔΔethA3ΔethA2: *ethA3*, *ethA2*, and *nudC* triple knockout strain; ΔΔethA3ΔethA1: *ethA3*, *ethA1* and *nudC* triple knockout strain.

To assess potential synergistic interactions between *MAB_0985* and *MAB_0103*, three triple knockout strains (ΔΔ0985Δ0103, ΔΔ3967Δ0985, and ΔΔ3967Δ0103) were generated from their corresponding double knockout parental strains (Figure S8). Simultaneous deletion of *MAB_0985* and *MAB_0103* in ΔnudC generated strain ΔΔ0985Δ0103, which exhibited high-level resistance to ETH, with a MIC restored to the WT level of 256 μg/mL (Table 2). This restoration indicates synergistic epistasis between *MAB_0985* and *MAB_0103*. The triple knockout strains ΔΔ3967Δ0985 and ΔΔ3967Δ0103 exhibited ETH susceptibility levels identical to those of the ΔΔ0985 and ΔΔ0103 strains, respectively (Table 2). This result aligns with our prior observation that deletion of *MAB_3967* does not further alter ETH susceptibility beyond the effects of deleting *MAB_0985* or *MAB_0103* (Table 2). DST results on solid agar media correlated with MIC values determined in liquid culture (Figure 3B). Furthermore, deletion of *MAB_0985*, *MAB_0103*, and *MAB_3967* in ΔnudC does not affect growth rate (Figure S3). No significant difference in growth rates was observed among the triple knockout strains ΔΔ0985Δ0103, ΔΔ3967Δ0985, and ΔΔ3967Δ0103. (Figure S3). Collectively, these data indicate that MAB_0985 and MAB_0103 are the most likely functional EthA homologs mediated ETH activation in *M. abscessus*.

### Overexpression of *MAB_0985*, *MAB_0103*, or *MAB_3967* in the ΔnudC and WT increased their susceptibility to ETH

The extra-chromosome plasmid pMV261 was used to overexpress *MAB_0985*, *MAB_0103*, or *MAB_3967* in ΔnudC. DST revealed that all strains exhibiting overexpression acquired sensitivity to ETH. The ETH susceptibility of all overexpression strains was enhanced, with MICs of 2 μg/mL to ΔnudC^C0985^ and nudC^C3967^, 1 μg/mL to ΔnudC^C0103^. These represent 8-, 8-, and 16-fold reductions in MIC relative to the parental strain, respectively (Table 2 and Table S2). Similarly, on solid agar plates, ΔnudC^C0985^, ΔnudC^C0103^, and ΔnudC^C3967^ were susceptible to ETH at 2 μg/mL, which is consistent with the liquid MIC results (Figure 4A). The growth rates of these overexpressed strains were not affected (Figure S3). To further validate their functions, we overexpressed *MAB_0985*, *MAB_0103*, and *MAB_3967* in the WT. The MIC of ETH to the WT^C0985^ was 128 μg/mL, representing a 2-fold decrease compared to the parental WT (Table 2). In contrast, no change in the MIC of ETH was observed for the WT^C0103^ and WT^C3967^. However, on agar plates, the WT^C0985^ exhibited sensitivity to ETH at 128 μg/mL, while the WT^C0103^ and WT^C3967^ displayed slight sensitivity at 256 μg/mL. This indicates that overexpression of *MAB_0103* and *MAB_3967* in the WT confers detectable increased susceptibility to ETH (Figure 4B). Quantitative reverse transcription-PCR (qRT-PCR) analysis demonstrated that, following 4-hours stimulation with 16 μg/mL ETH, the mRNA level of *MAB_0985* in ΔnudC increased 5.65-fold, whereas *MAB_0103* expression exhibited a modest 1.40-fold increase (Figure 4C). Conversely, *MAB_3967* transcript levels exhibited no significant upregulation (Figure 4C). Analysis of drug sensitivity profiles from the ΔnudC^C3967^, Δ3967, ΔΔ3967, ΔΔ0985Δ3967, and ΔΔ0103Δ3967 strains, we speculate that MAB_3967 (EthA3) may also function as a potential EthA in *M. abscessus*. Its role in ETH activation may only become detectable under conditions of substantial overexpression.

**Figure 4.**
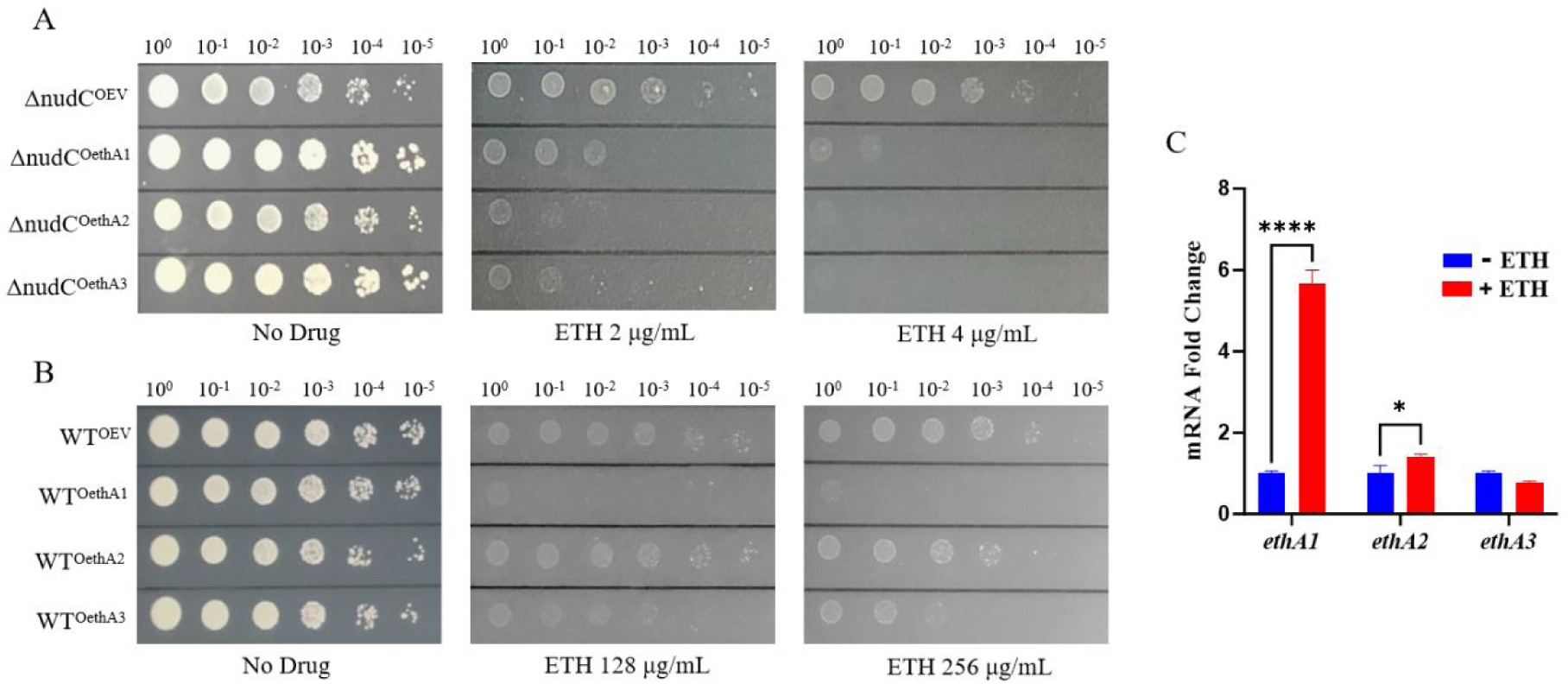
Overexpression of *ethA1* (*MAB_0985*)*, ethA2* (*MAB_0103*), or *ethA3* (*MAB_3967*) in the *nudC* knockout strain or wild-type *M. abscessus* enhances the susceptibility to ETH. (A) Susceptibility of *nudC* knockout strains overexpressing *ethA1*, *ethA2,* or *ethA3* to ETH. ΔnudC^OEV^, ΔnudC^OethA^^1^, ΔnudC^OethA^^2^, ΔnudC^OethA^^3^: *M. abscessus nudC* knockout strain containing an empty vector or the vector for overexpressing *ethA1*^Mab^, *ethA2*^Mab^, or *ethA3*^Mab^, respectively. (B) Susceptibility of wild-type *M. abscessus* overexpressing *ethA1*, *ethA2,* or *ethA3* to ETH. WT^OEV^, WT^OethA^^1^, WT^OethA^^2^, WT^OethA^^3^: wild-type *M. abscessus* containing an empty vector or the vector for overexpressing *ethA1*^Mab^, *ethA2*^Mab^, or *ethA3*^Mab^, respectively. (C) Gene expression levels of *ethA1*, *ethA2*, and *ethA3* in the *nudC* knockout strain in the presence and absence of ETH. \**P* < 0.05, \*\*\*\**P* < 0.0001. Data are means ± SD of three biological duplicates.

### Deletion of *MAB_0985* or *MAB_0103* reduces the formation of the ETH-NAD adduct in ΔnudC treated with ETH

Our prior research demonstrated that *nudC* deletion in the WT resulted in the accumulation of an ETH-NAD adduct, which significantly increased susceptibility to ETH. Therefore, we employed high-performance liquid chromatography–mass spectrometry (HPLC-MS) to quantify ETH-NAD adduct levels in the double and triple knockout strains to determine the contribution of MAB_0985 and MAB_0103 to ETH activation. HPLC-MS analysis revealed that under 256 μg/mL ETH treatment, the levels of ETH-NAD adduct were significantly reduced in both double knockout strains (ΔΔ0985 and ΔΔ0103), with the triple knockout strain ΔΔ0985Δ0103 exhibiting an even more pronounced reduction (Figure 1C). In contrast, deletion of *MAB_3967* in ΔnudC, ΔΔ0985, or ΔΔ0103 did not alter ETH-NAD adduct production. These findings align with the DST results, confirming that MAB_0985 (EthA1) and MAB_0103 (EthA2) serve as the principal EthA enzymes in *M. abscessus*.

### EthR inhibits *ethA1* expression but does not influence *ethA2* and *ethA3* expression

Intriguingly, *ethR* (*MAB_0984c*) is located adjacent to the *ethA1* (*MAB_0985*) locus. The genetic organization of the *MAB_0984c*-*MAB_0985* locus is identical to that of the Mtb *ethR*-*ethA* (*Rv3854c*-*Rv3855*) locus, with both comprising adjacent open reading frames separated by short intergenic regions (IGR, 75 bp and 74 bp, respectively) (Figure 5A). Structural homology with the Mtb *ethA*-*ethR* regulon indicates that EthR acts as a repressor of *ethA1* expression, likely by modulating the activity of its IGR. Bioinformatic analysis of the IGR between *ethR* and *ethA1* identified consensus -10 and -35 promoter elements, confirming that this IGR functions as the promoter for *ethA1* (Figure S9). IGR-enhanced green fluorescent protein (eGFP) reporter strains were engineered to monitor EthR-mediated suppression of *ethA1*. Quantitative analysis of reporter strains in the WT demonstrated that fluorescence signals from *MAB_0985-*, *MAB_0103-*, and *MAB_3967*-IGR-eGFP fusion constructs were markedly less intense than those elicited by the *hsp60*-eGFP positive control. However, overexpression of the *ethA1*-IGR-eGFP reporter in the ΔethR mutant increased fluorescence by more than 27-fold relative to WT levels, producing a signal exceeding that of the *hsp60*-eGFP positive control (Figure 5B,C). Conversely, no significant change in eGFP fluorescence intensity was observed for *MAB_0103*-IGR and *MAB_3967*-IGR promoters in the ΔethR mutant compared to the WT (Figure 5B, C). This indicates that expression from these promoters is not regulated by EthR. Following *ethR* deletion in the ΔnudC strain, *ethA1* transcript levels increased over 300-fold (quantified by qRT-PCR), whereas *ethA2* and *ethA3* expression remained unchanged (Figure 5D). Following treatment with 0.125 μg/mL ETH, *ethA1* mRNA level in the ΔΔethR increased over 700-fold, significantly exceeding the minimal changes observed for *MAB_0103* (3.37-fold) and *MAB_3967* (1.96-fold) (Figure 5E). These findings demonstrate that EthR specifically suppresses *ethA1* expression, while leaving *ethA2* and *ethA3* expression unaffected. Accordingly, *ethR* deletion significantly upregulated *ethA1* expression.

**Figure 5.**
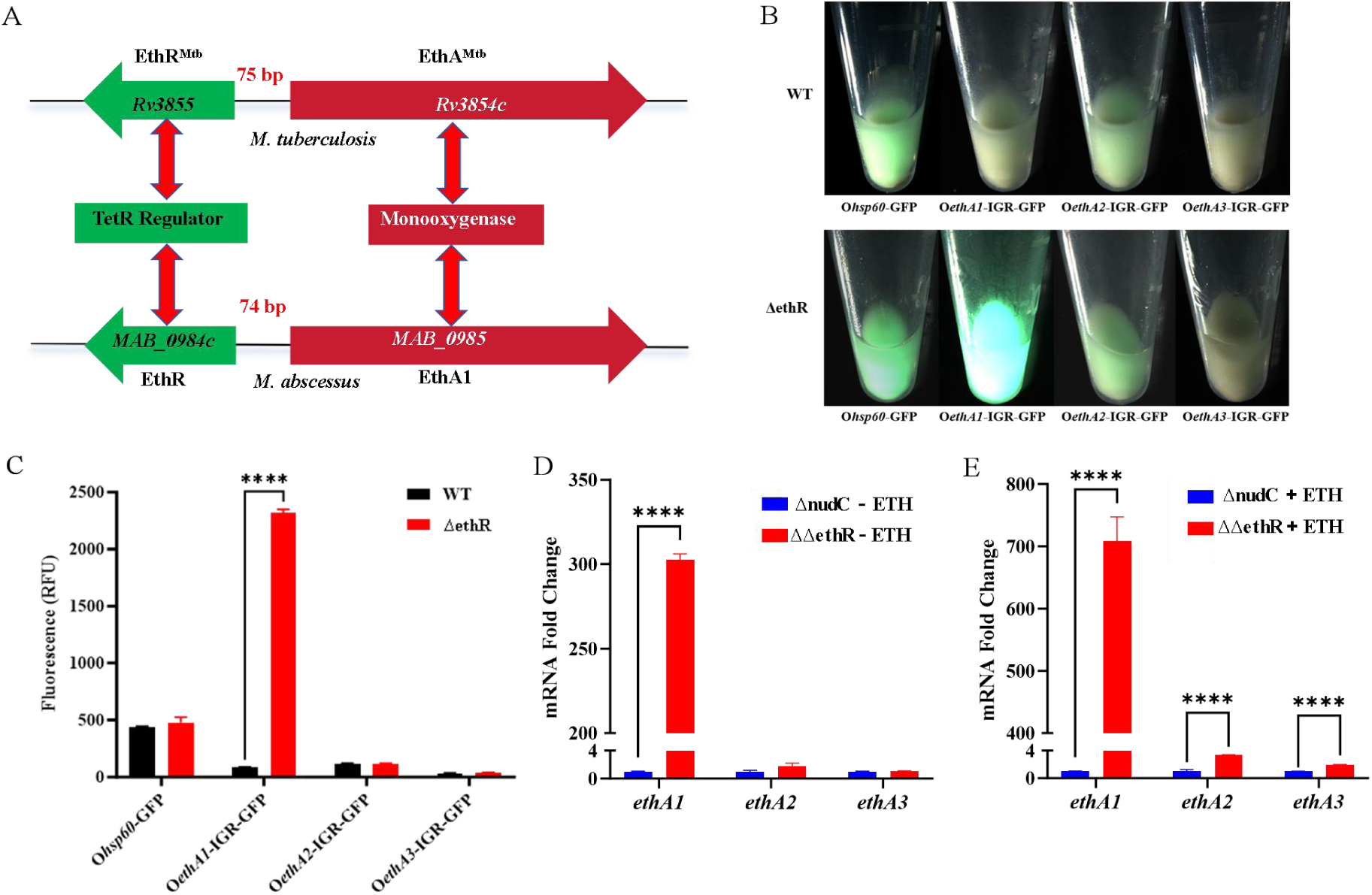
EthR specifically suppresses *ethA1* expression but does not influence *ethA2* and *ethA3* expression. (A) Comparison of the genetic organization and predicted protein functions at homologous loci between *M. abscessus* and *M. tuberculosis*. Genes *MAB_0984c* and *MAB_0985* correspond to *ethR*^Mtb^ (*Rv3855*) and *ethA*^Mtb^ (*Rv3854c*), respectively. (B-C) Overexpressing *ethA1*-intergenic region (IGR)-GFP in *ethR* knockout strain, resulting in a significant increase in relative fluorescence (RFU). WT: wild-type *M. abscessus*; ΔethR: *ethR* knockout strain; O*hsp60*-GFP, O*ethA1*-IGR-GFP, O*ethA2*-IGR-GFP, O*ethA3*-IGR-GFP: overexpressing *hsp60*-GFP, *ethA1*(*MAB_0985*)-IGR-GFP, *ethA2* (*MAB_0103*)-IGR-GFP, *ethA3* (*MAB_3967*)-IGR-GFP in WT and ΔethR, respectively. (D-E) Gene expression levels of *ethA1*, *ethA2*, and *ethA3* in the *ethR* and *nudC* double knockout strain in the absence and presence of ETH. ΔnudC: *M. abscessus nudC* knockout strain; ΔΔethR: *ethR* and *nudC* double knockout strain; \*\*\*\**P* < 0.0001. Data are means ± SD of three biological duplicates.

### Molecular docking predicts high-affinity binding of BDM31343 to the predicted EthR structure

The three-dimensional structure of EthR was computationally predicted using AlphaFold3 and validated via a per-residue confidence score (pLDDT) of 88.4, indicating high model reliability (Figure 6A). Moreover, the predicted structure of EthR exhibits high homology to that of EthR^Mtb^ (PDB code: 3TP0), with both proteins containing nine alpha-helices (Figure 6A). BDM31343 is an inhibitor of EthR^Mtb^, which increases the antibacterial efficacy of ETH against *M. tuberculosis* by 10-fold^14^. To elucidate the potential binding of BDM31343 to EthR, molecular docking was performed to predict the binding affinity and specific interaction modes. The molecular docking result shows that the inhibitor BDM31343 fits well into the binding pocket of the EthR and is stabilized by multiple non-covalent interactions. The predicted binding free energy for the BDM31343-EthR complex is -10.54 kcal/mol. The binding is anchored by a specific hydrogen bond with ASN-183 (Figure 6B). This conformation is further stabilized by strong π-π stacking interactions with PHE-114 and hydrophobic contacts involving residues such as TRP-142, ILE-152, and LEU-91 (Figure 6B). This synergistic interplay of interactions underlies the potent and specific binding affinity of BDM31343, enabling its effective inhibition of the EthR protein.

**Figure 6.**
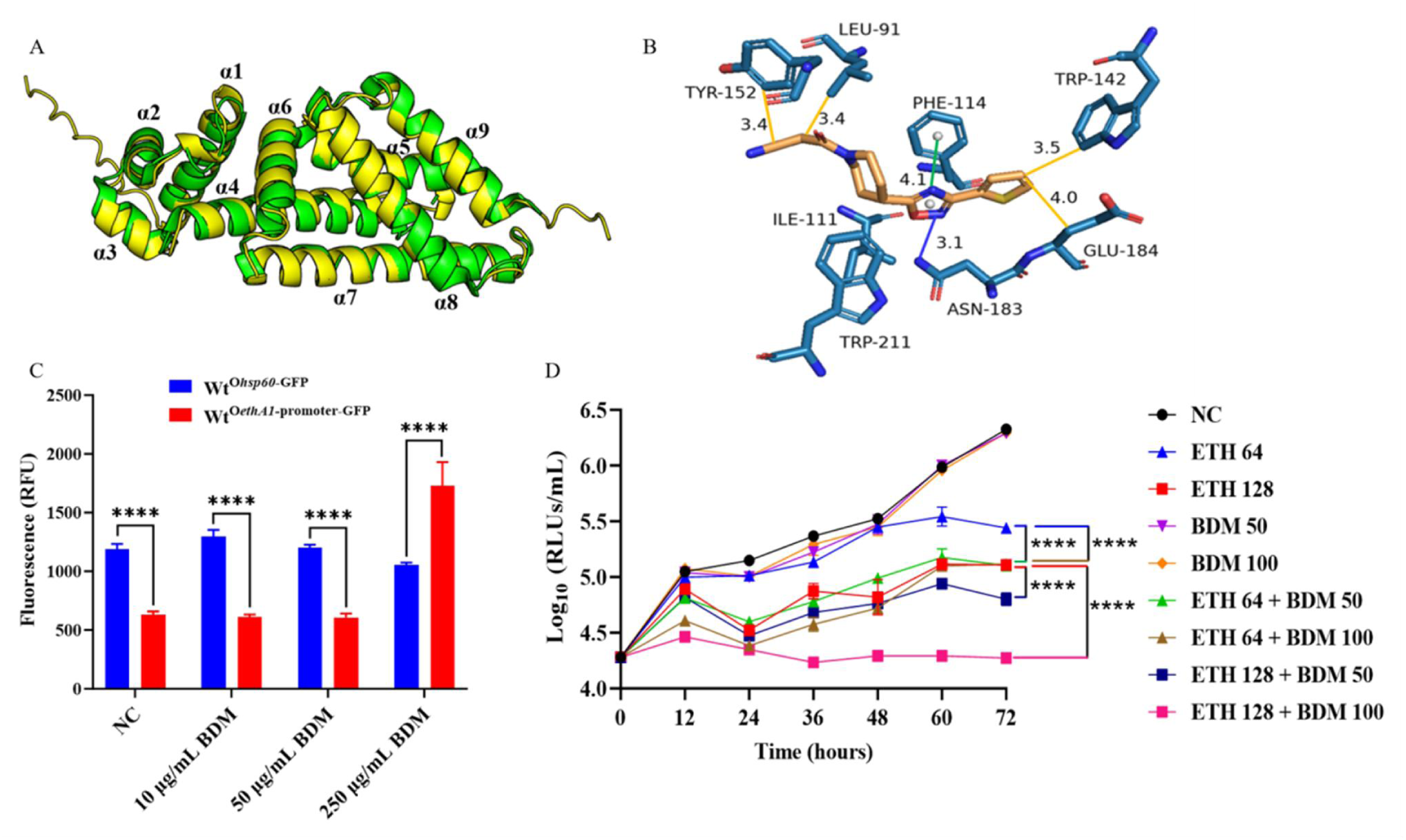
EthR^Mtb^ inhibitor BDM31343 can boost the efficacy of ETH against *M. abscessus* via suppressesion of EthR. (A) The structure of EThR in *M. abscessus* is highly homologous to that of EthR^Mtb^ (PDB code: 3TP0). Green: EthR^Mtb^ structure; Yellow: *M. abscessus* EThR structure predicted by AlphaFold3. (B) Interaction patterns between BDM31343 and the amino acid residues of EthR. Blue line: hydrogen bond; Green line: π-π stacking interaction; Yellow line: hydrophobic interaction. (C) High doses of BDM31343 (BDM) resulted in a significant increase in the relative fluorescence (RFU) in wild-type *M. abscessus* overexpressing *ethA1*-promoter-GFP. WT^O*hsp*60-GFP^, WT^O*ethA*1-promoter-GFP^: wild-type *M. abscessus* overexpressing *hsp60*-GFP and *ethA1*-promoter-GFP, respectively. NC: negative control. \*\*\*\**P* < 0.0001. (D) Time-kill curves of BDM31343 and ETH against AlMab. RLUs: relative light units. Numbers following the ETH and BDM indicate the working concentrations (µg/mL). \*\*\*\**P* < 0.0001.

### BDM31343 potentiates ETH activity against *M. abscessus* via inhibition of EthR

To further evaluate BDM31343’s inhibitory effect on EthR, the compound was administered to cultures of the IGR-eGFP reporter strain. When the concentration of BDM31343 reached 250 μg/mL, a marked increase in the *ethA1* promoter activity was observed, resulting in the eGFP fluorescence intensity surpassing even that of the *hsp60*-eGFP control (Figure 6C). To further validate the efficacy of BDM31343, we employed an autoluminescent *M. abscessus* (AlMab)^17^ strain to assess whether this compound potentiates the antibacterial activity of ETH against *M. abscessus*. Both concentrations of BDM31343 (50 and 100 μg/mL) produced a 0.33 log_10_ relative light units (RLUs) decrease in AlMab under 64 μg/mL ETH, with growth inhibition equivalent to 128 μg/mL ETH treatment (Figure 6D). At 128 μg/mL ETH, the reduction in AlMab RLUs was greater at 100 μg/mL BDM31343 (0.83 log_10_) than at 50 μg/mL (0.31 log_10_), while complete growth inhibition occurred only at the higher concentration (Figure 6D). These findings demonstrate that high-dose BDM31343 inhibits EthR and enhances *M. abscessus* susceptibility to ETH, albeit with reduced efficacy compared to its activity against *M. tuberculosis*. Therefore, developing small-molecule inhibitors targeting *M. abscessus* EthR represents a promising strategy to enhance the antibacterial efficacy of ETH.

## DISCUSSION

*M. abscessus* is a predominant rapidly growing NTM causing both pulmonary and extrapulmonary infections. The intrinsic resistance of *M. abscessus* to antibiotics, attributable to its unique cell wall architecture and metabolic mechanisms, constitutes a formidable challenge in clinical management^18^. Therefore, elucidating the molecular mechanisms of drug resistance in *M. abscessus* is critically important for clinical management. Given this bacterium’s unique resistance profile, and poor treatment outcomes, the identification of novel drug targets and resistance-modulating strategies remains an urgent priority.

ETH is one of the most effective second-line drugs for the treatment of multidrug-resistant *M. tuberculosis*. However, its clinical utility is limited by dose-dependent adverse effects, including gastrointestinal toxicity, which often compromise treatment adherence^14^. Strategies that enable ETH dose reduction while preserving antibacterial efficacy are therefore highly desirable. In *M. tuberculosis*, ETH is activated by the Baeyer-Villiger monooxygenase (BVMO) EthA^10,19,20^, resulting in a stable ETH-NAD adduct, which subsequently inhibits the InhA, thereby disrupting mycolic acid biosynthesis^21,22^. Since the expression of EthA is regulated by the TetR-family transcriptional repressor EthR^23^, small-molecule inhibitors of EthR have been successfully developed for *M. tuberculosis* to stimulate EthA expression^14,24–27^, thereby potentiating ETH activation and enhancing its antibacterial activity^13,14^. Although EthR and EthA are well-characterized in *M. tuberculosis*, their homologous counterparts and associated functions in *M. abscessus* are yet to be elucidated. In this study, we identify EthR and its cognate target EthA1 in *M. abscessus* for the first time. Furthermore, we demonstrated that disrupting this regulatory mechanism increases susceptibility to ETH.

It is well established that homologous proteins with amino acid sequence identity exceeding 40% are highly likely to share functional similarity^28^. BLASTp analysis using the EthA^Mtb^ sequence identified three putative homologs in *M. abscessus*: MAB_0985, MAB_0103, and MAB_3967, which we designated as EthA1, EthA2, and EthA3, respectively. Each shares over 40% amino acid sequence identity with EthA^Mtb^. EthA1 and EthA2 are the primary functional EthAs in *M. abscessus*. Disruption of either gene in ΔnudC significantly reduced ETH-NAD adduct formation and consequently enhanced bacterial resistance to ETH. Remarkably, the combined deletion of *ethA1* and *ethA2* in the ΔnudC abolished ETH activation in the resulting triple mutant. This reduction in ETH-NAD adduct formation conferred high-level ETH resistance. Conversely, overexpression of either *ethA1* or *ethA1* in the ΔnudC enhanced susceptibility to ETH.Following treatment with ETH, the ΔnudC mutant exhibited a significant upregulation of *ethA1* transcription and a modest increase in *ethA2* expression. Conversely, deletion of *ethA3* did not significantly affect susceptibility to ETH in either the wild-type or ΔnudC strain.No significant change in *ethA3* expression was observed in the ΔnudC following ETH treatment. Collectively, these data imply that EthA3 does not appear to be a major EthA in *M. abscessus*. However, overexpression of *ethA3* in the ΔnudC using the multicopy plasmid pMV261 conferred high susceptibility to ETH. Therefore, EthA3 should be regarded as a functional EthA in *M. abscessus*, yet its physiological role appears minimal under native conditions. Functional significance emerges only upon overexpression, likely due to its inherently low catalytic efficiency. Therefore, EthA1 exhibits the strongest ETH-activating capability, followed by EthA2, while EthA3 is the weakest (Figure 7). Moreover, the conserved BVMO motif [FXGXXXHXXXW(P/D)]^29^, characteristic of EthA in *M. tuberculosis*, was identified in EthA1, EthA2, and EthA3 of *M. abscessus* (Figure S4). Under physiological conditions, BMVOs are primarily involved in oxidative degradation of endogenous substrates such as ketones and steroids^30^. The differential activation efficiency of ETH by EthA1, EthA2, and EthA3 likely arises from their distinct substrate specificities, as each BVMO typically exhibits selectivity toward specific compound classes^12^, and ETH is not a natural substrate for BMVOs. Like *ethA1*, the *ethA2* and *ethA3* genes are likely regulated by their cognate EthR protein. However, the genes adjacent to *ethA2* and *ethA3* do not belong to either the TetR family of repressors or the AraC/XylS family of transcriptional factors, indicating that their specific transcriptional regulators remain to be identified.

**Figure 7.**
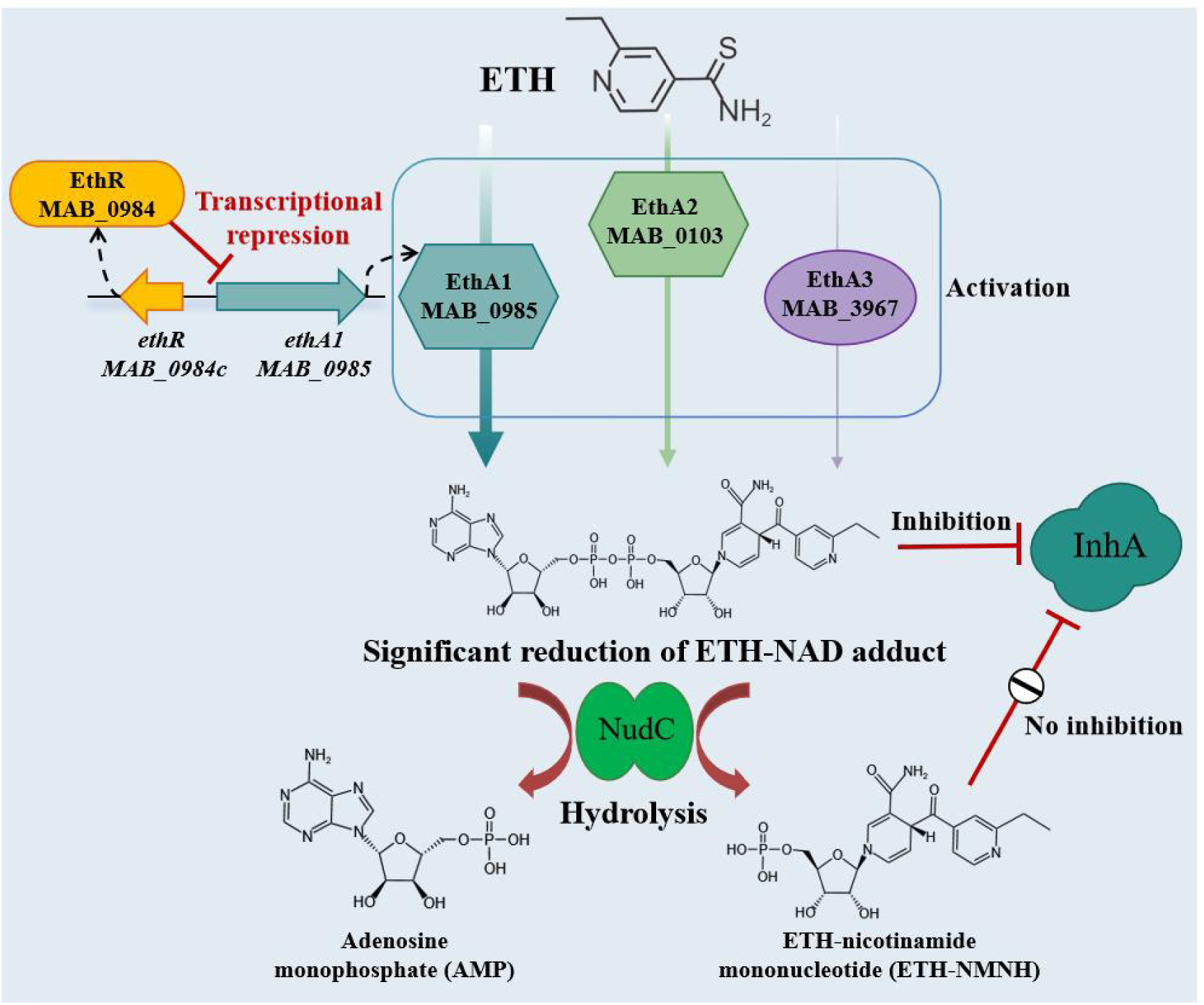
Schematic representation of the resistance mechanism of *M. abscessus* to ETH. ETH is activated by EthA1 (MAB_0985), EthA2 (MAB_0103), and EthA3 (MAB_3967) to form the ETH-NAD adduct in *M. abscessus.* Among them, EthA1 exhibits the strongest activating potency, followed by EthA2, while EthA3 is the weakest. EthR specifically suppresses *ethA1* expression, resulting significant reduction of ETH-NAD adduct in *M. abscessus.* The ETH-NAD adduct can be hydrolyzed by NudC into adenosine monophosphate (AMP) and ETH-nicotinamide mononucleotide (ETH-NMNH). This reaction eliminates the adduct’s inhibition of InhA, consequently conferring resistance to ETH in *M.abscessus*. The thickness of the lines connected to ETH represents the ETH-activating capability.

Our previous study revealed a mechanism of ETH resistance in *M. abscessus* mediated by NudC, encoded by *MAB_3513c*^15^. NudC functions as a pyrophosphohydrolase that hydrolyzes the ETH-NAD adduct, thereby counteracting ETH toxicity. The proline residue at position 226 was shown to be critical for NudC dimerization and hydrolase activity, which underlies the intrinsic ETH resistance. However, the *nudC* knockout reduced the MIC of ETH against *M. abscessus* from 256 μg/mL to 16 μg/mL, which was still higher than the MIC of ETH for *M. tuberculosis*, suggesting the existence of additional resistance mechanisms in *M. abscessus*. In *M. tuberculosis*, CinA has been shown to confer cross-tolerance to INH, ETH, delamanid, and pretomanid by hydrolyzing drug-NAD adducts, functioning in a manner similar to NudC^31^. Whether a CinA homolog in *M. abscessus* contributes to ETH resistance, in addition to NudC, has not been reported. However, knockout of *MAB_2443*, the putative *cinA* homolog, in either the WT or ΔnudC strain did not affect *M. abscessus* susceptibility to ETH or INH (Table 2). These results indicate that CinA^Mab^ does not fulfill the same functional role as CinA^Mtb^, and the underlying mechanism warrants further investigation.

In this study, we identified MAB_0984 as an additional key determinant of ETH resistance in *M. abscessus*, which functions as EthR by inhibiting the expression of *ethA1* to reduce ETH activation. Bioinformatic analysis revealed that *ethA1* is adjacent to the TetR-family transcriptional regulator gene *MAB_0984c*, mirroring the genomic organization of the *ethR-ethA* locus (*Rv3854c*-*Rv3855*) in *M. tuberculosis*. MAB_0984 exhibits significant sequence identity with *M. tuberculosis* EthR, strongly supporting its classification as a functional homolog of this transcriptional regulator. The increased susceptibility to ETH observed upon *ethR* knockout was substantially greater than that resulting from *nudC* deletion, indicating EthR’s more critical role in conferring intrinsic resistance. Concurrent disruption of *nudC* and *ethR* produced a synergistic effect, reducing the MIC substantially below that observed with either single knockout. Metabolite analysis directly links the observed phenotypic differences to underlying mechanistic changes. While *nudC* disruption impairs adduct clearance, leading to moderate accumulation, the significantly greater buildup in the ΔΔethR mutant reflects profound dysregulation of the ETH activation pathway. These findings demonstrate that intrinsic ETH resistance in *M. abscessus* is primarily governed by the EthR-regulated activation pathway rather than the NudC-mediated clearance pathway. Disrupting EthR substantially enhances susceptibility to ETH, establishing it as the primary drug target. However, complete reversal of resistance necessitates dual targeting of both pathways, as inhibition of either mechanism alone is insufficient to overcome this robust intrinsic barrier. Mechanistic studies using qRT-PCR and an IGR-eGFP reporter system confirmed that EthR inhibits *ethA1* expression by regulating the activity of the promoter. Collectively, these results demonstrate that MAB_0984 functions as the EthR regulator in *M. abscessus*, which inhibits EthA1 production and thereby mediates intrinsic resistance to ETH (Figure 7).

Small-molecule compounds targeting EthR^Mtb^ have been found to significantly boost the efficacy of ETH both *in vitro* and *in vivo*. Co-administration of the EthR-targeting compounds BDM31343 or BDM41906 with ETH dramatically lowers the required ETH dose without compromising efficacy, achieving mycobacterial load comparable to standard high-dose treatment^14,27^. However, they failed to enhance the activity of ETH against *ethA*-mutant strains. Subsequently, EthA2 and EthR2 were identified in *M. tuberculosis*, prompting the development of SMARt420, a specific inhibitor targeting EthR2^11^. SMARt420 not only completely reversed acquired ETH resistance and eradicated ETH-resistant bacterial loads in mice, but also enhanced the baseline susceptibility of *M. tuberculosis* to ETH. The recently identified compound SMARt751 targets the *virS* transcriptional regulator in *M. tuberculosis* to modulate the *mymA* operon, which encodes a monooxygenase capable of activating ETH^13^. SMARt751 significantly boosted the therapeutic efficacy of ETH in both *in vitro* assays and murine models of acute/chronic tuberculosis. Moreover, it restored the full therapeutic efficacy of ETH in mice infected with ETH-resistant strains. SMARt751 uniquely circumvents the predominant resistance mechanisms found in clinical *M. tuberculosis* isolates, such as alterations in the *ethA, ethR* promoter, and *inhA* promoter, thereby overcoming typical forms of ETH resistance. However, no homologs of *ethR2* or *virS* could be identified in the *M. abscessus*. The EthR^Mtb^ inhibitor BDM31343 exhibits cross-species activity by inhibiting EthR^Mab^, as shown by increased eGFP signal in *M. abscessus* containing an *ethA1-*promoter-eGFP construct. BDM31343 enhances the efficacy of ETH against *M. abscessus*, albeit at elevated concentrations. Thus, developing small-molecule inhibitors specifically targeting EthR represents a promising strategy to effectively potentiate ETH-based treatment in *M. abscessus*.

In conclusion, this study identifies MAB_0984 as the transcriptional repressor EthR, which specifically suppresses *ethA1* expression, resulting significant reduction of the ETH-NAD adduct in *M. abscessus.* The intrinsic resistance of *M. abscessus* to ETH is primarily governed by the EthR-regulated activation pathway rather than the NudC-mediated clearance pathway. We identify EthA1 (MAB_0985) as the most potent activator of ETH, with EthA2 (MAB_0103) exhibiting intermediate activity and EthA3 (MAB_3967) demonstrating the weakest activation. The development of EthR inhibitors represents a promising avenue to sensitize mycobacteria to ETH. This work elucidates the mechanism of ETH resistance in Mab, establishing a conceptual and mechanistic framework for developing more effective ETH-based therapies.

## MATERIALS AND METHODS

### Bacterial strains and culture conditions

*E. coli* Trans T1 (Beijing TransGen Biotech) was used for plasmid propagation and cultured at 37°C in liquid or solid LB medium. The strain *M. abscessus* GZ002 (NCBI GenBank accession number CP034181) isolated and characterized in our laboratory, was used as the parental strain throughout this study. It was cultured at 37°C in 7H9 broth (containing 2% glycerol, 10% OADC, and 0.05% Tween 80) or on 7H10 agar (supplemented with 2% glycerol and 10% OADC). When necessary, kanamycin (Kan, 50 μg/mL) or zeocin (Zeo, 30 μg/mL) was added to the medium for plasmid selection and maintenance. Anhydrotetracycline (ATc) was used as an inducer at a concentration of 100 ng/mL. The autoluminescent *M. abscessus* strain (AlMab), engineered in our laboratory, was used for drug susceptibility testing based on luminescence readouts.

### Identification, knockout and complementation of putative *ethR* and *ethAs* homologs

Putative *ethR* and *ethA* homologs in *M. abscessus* were identified by BLASTp analysis using characterized EthR and EthA proteins from *M. tuberculosis* as query sequences. Genes exhibiting > 40% amino acid sequence identity were selected for functional analysis.

Gene knockouts were generated using a CRISPR-NHEJ system. Briefly, the temperature-sensitive plasmid pCR-Zeo (generously provided by Prof. Yicheng Sun’s laboratory) was utilized for crRNA expression to generate knockout mutants. Specifically, primers harboring the gene-specific crRNA sequence were designed, annealed, and ligated into the digested pCR-Zeo backbone. Recombinant plasmids were introduced into *M. abscessus* harboring the pNHEJ-cpf1 plasmid by electroporation.

Transformants were plated on Middlebrook 7H11 agar supplemented with Kan (100 μg/mL), Zeo (30 μg/mL), and ATc (200 ng/mL), and incubated at 30℃ for 72 hours. Single colonies were screened by PCR, followed by agarose gel electrophoresis and Sanger sequencing (performed by Sangon Biotech, Shanghai) to confirm the construction of the desired knockout strains. Complementation strains were constructed by introducing the respective wild-type genes cloned into the integrative plasmid pMV261.

### DST

The MICs were determined by the broth microdilution method. WT and the putative *ethA* and *ethR* knockout and complemented strains were first grown in 7H9 medium to the mid-log phase (OD₆₀₀ = 0.6-0.8). The cultures were then diluted to an OD₆₀₀ of 0.1-0.2 in Tween-80-free 7H9 medium, and subsequently diluted 1000-fold to prepare the final inoculum for testing. The drugs were subjected to twofold serial dilution. The assay was performed with four technical replicates for each concentration in three independent biological replicates. We performed DST on solid medium by spotting ten-fold serial dilutions (up to 10⁻⁵) of mid-log phase bacterial cultures onto 7H10 agar plates containing specified drug concentrations. Antibiotic-free plates were used as negative controls, and all assays were repeated three times.

### Determination of growth curves

Bacterial growth was monitored by measuring the absorbance at 600 nm. Mid-exponential phase cultures of *M. abscessus* were diluted to an OD_600_ of 0.05 and incubated at 37℃ with shaking at 200 rpm. The OD_600_ was measured at 6-hour intervals for a total duration of 72 hours using a spectrophotometer. Experiments were performed in triplicate, and the growth curves were plotted using GraphPad Prism 10.1.2.

### Extraction of ETH metabolites

ETH metabolites were extracted as previously described^15^, with minor modifications. Briefly, bacterial strains were cultured to an OD6_00_ of 0.7 and then exposed to 256 μg/mL ETH for 3 hours at 37°C in a shaking incubator. Bacteria were harvested and washed twice with deionized water by centrifugation at 13,000 × g for 10 minutes at 4°C. The obtained bacterial precipitate was resuspended in 6 mL of chloroform-methanol (2:1), incubated at room temperature with shaking at 200 rpm for 3 hours, followed by solvent evaporation using a CentriVap concentrator (Labconco). The dried bacterial pellet was resuspended in 100 mM ammonium carbonate (pH 7.0) and subjected to vigorous mixing with 0.1 mm and 0.5 mm zirconia beads for physical disruption. The lysate was centrifuged at 18,000 × g for 20 minutes at 4°C, and the resulting supernatant was collected and passed through a 0.22 μM filter (Millipore) to remove cellular debris and large particles. The final filtrate was acidified with formic acid to a final concentration of 0.1% (v/v), followed by centrifugation at 18,000 × g for 20 minutes at 4°C. Subsequently, a 2 μL aliquot of the each resulting product was analyzed by HPLC-MS using an Ultimate 3000/Q-Exactive HF-X system (ThermoFisher Scientific).

### Detection of ETH-NAD adducts

HPLC-MS analysis was employed to detect the ETH-NAD adduct. Chromatographic separation was performed by injecting 2 μL of sample onto an ACQUITY UPLC BEH C18 column (100 × 2.1 mm, 1.7 μm; Waters). Chromatographic separation was performed at a flow rate of 0.3 mL/min using a linear gradient of 0.1% formic acid in water (mobile phase A) and acetonitrile (mobile phase B): 1-90% B over 4 min, followed by 90% B for 1 min and re-equilibration at 1% B for 2 min.. MS detection was conducted on an Ultimate 3000/Q Exactive HF-X system (Thermo Fisher Scientific) in positive ion mode over an m/z range of 100-1000 withworking parameters as follows: sheath gas flow rate: 40, auxiliary gas flow rate: 10, spray voltage: 3.5kV, capillary temperature: 300℃, S-lens RF level: 55, and auxiliary gas heater temperature: 350℃.

### Construction of eGFP reporter strains and qualitative/quantitative fluorescence analysis

To construct the reporter plasmids, the IGR of *ethA1*, *ethA2*, and *ethA3* were amplified and cloned upstream of the *egfp* gene in the integrative plasmid pRH2502. These plasmids were then electroporated into WT and ΔethR strains. The transformation mixture was plated on 7H10 agar plates containing Kan (100 μg/mL) and incubated at 37°C for 72 hours. Single colonies were then selected for PCR amplification and subsequent DNA sequencing to confirm the correct constructs, resulting in the corresponding eGFP reporter strains. For qualitative fluorescence analysis, cells from a 10 mL culture of the eGFP reporter strain (OD₆₀₀ = 0.8) were harvested and visualized using a fluorescence microscope (Leica M205 FA). For quantitative analysis, 150 μL aliquots of exponentially growing cultures of the eGFP reporter strain were dispensed in triplicate into white 96-well plates and incubated at 37°C for 72 hours. eGFP fluorescence (excitation 488 nm, emission 520 nm) was measured using a FlexStation 3 microplate reader (Molecular Devices).

### RNA extraction and qRT-PCR

ΔnudC and ΔΔethR strains were grown to mid-log phase and treated with ETH at their respective MICs (16 and 0.125 μg/mL) for 4 hours, with equal volumes of DMSO serving as the negative controls for each treatment. Total RNA was extracted using the HiPure Bacterial RNA Kit (Magen) and quantified via Nanodrop UV spectrophotometry (Thermo Fisher Scientific). RT-qPCR were conducted using the HiScript III All-in-one RT SuperMix Perfect for qPCR (Vazyme) and Taq Pro Universal SYBR qPCR Master Mix (Vazyme) on a CFX96 Touch Real-Time PCR Detection System (Bio-Rad). Quantitative PCR data were analyzed using the 2^-ΔΔ*C*t^ method, with relative mRNA expression of target genes normalized to the reference gene *sigA*^32^.

### Molecular docking analysis

AlphaFold3 (https://alphafoldserver.com/) was used to predict the structure of the EthR. The structure of BDM31343 was obtained from the NCBI PubChem database (https://pubchem.ncbi.nlm.nih.gov/). Structural alignment of *M. abscessus* EthR with its homologous protein EthR^Mtb^ (PDB code: 3TP0) was performed using PyMOL (http://www.pymol.org). AutoDock Tools version 1.5.6 was employed to predict the binding affinity and interaction modes between BDM31343 and the predicted structure of EthR.

### Evaluation of BDM31343 on ETH susceptibility using the AlMab

The AlMab culture was grown to a density of 1 × 10⁶ RLUs, and then adjusted to a working concentration of 3,000-5,000 RLUs. In 1.5 mL microcentrifuge tubes, 5 μL of BDM31343/ETH combinations (in DMSO) were mixed with 195 μL of diluted AlMab suspension to a final volume of 200 μL, followed by mixing and incubation at 37°C. For the negative control, 5 μL of DMSO was combined with 195 μL of AlMab. RLUs values were monitored every 12 hours with a GloMax 20/20 luminometer during the 72-hour incubation period.

### Statistical analysis

Data were analyzed with GraphPad Prism 10.1.2 (GraphPad software). Statistical significance was assessed using unpaired Student’s *t*-test or one-way ANOVA, with statistical significance set at *P* < 0.05.

## ACKNOWLEDGMENTS

This study was supported by the National Key R&D Program of China (2021YFA1300904), the Major Project of Guangzhou National Laboratory (GZNL2025C01003), the National Natural Science Foundation of China (82302554 and 82502762), the China-Maurice Wilkins Centre (MWC) Programme, Guangdong-New Zealand Bilateral Joint Funding (2025A0505020001), the Key Research and Development Program of Guangzhou (2025B01J3019) and the Guangzhou National Laboratory and State Key Laboratory of Respiratory Disease (GZNL2025B01006). We also extend our sincere appreciation to Cuicui Liu of the Chinese Academy of Sciences for her assistance in detecting the ETH-NAD adduct across various *M. abscessus* strains.

## AUTHOR CONTRIBUTIONS

**Xiaofan Zhang**: Conceptualization (equal); methodology (lead); investigation (lead); data curation (lead); formal analysis (lead); validation (lead); funding acquisition (lead); writing-original draft (lead); writing-review and editing (lead). **Lijie Li**: Investigation (equal) and validation (equal). **Ziwen Lu**: Investigation (equal) and software (equal). **Zhaohui Duan**: Methodology (supporting) and formal analysis (equal). **H.M. Adnan Hameed**: Writing-original draft (supporting) and Writing-review (lead). **Belachew Aweke Mulu**: Formal analysis (equal) and software (supporting). **Cuiting Fang**: Investigation (supporting) and validation (supporting). **Xirong Tian**: Investigation (supporting) and validation (supporting). **Xinyue Wang**: Formal analysis (supporting) and validation (supporting). **Hongyi Chen**: Validation (supporting) and writing-review (supporting). **Liqiang Feng**: Conceptualization (supporting) and writing-review (supporting). **Matthew B. McNei**l: Formal analysis (supporting) and writing-review (supporting). **Xindong Liu**: Formal analysis (supporting) and writing-review (supporting). **Shuai Wang**: Conceptualization (equal); project administration (lead); funding acquisition (lead); writing-review and editing (lead). **Tianyu Zhang**: Conceptualization (equal); project administration (lead); funding acquisition (lead); writing-review and editing (lead).

## ETHICS STATEMENT

This work did not involve human participants or animal experiments, and thus no ethical review, approval, or consent was required.

## DATA AVAILABILITY

The authors declare that all data supporting the findings of this study are provided within the article and its supplementary materials.

