## Supplemental Material for "Identification and characterization of *ethR* and *ethA* genes impacting the sensitivity of *Mycobacterium abscessus* to ethionamide"

**Table S1. The susceptibility of *M. abscessus ethA1* or *ethA2* knockout strains in a *nudC* deletion background, as well as their complementary strains, to ETH and INH.**

| ***M. abscessus* strains** | **Antibiotics/ MICs[a] (μg/mL)** | |
|---|---|---|
| | ETH | INH |
| WT | 256 | > 512 |
| ΔnudC | 16 | 1 |
| ΔΔethA1 | 64 | 1 |
| ΔΔethA1$^{CMab}$ | 16 | 1 |
| ΔΔethA1$^{CMtb}$ | 32 | 1 |
| ΔΔethA1$^{CEV}$ | 64 | 1 |
| ΔΔethA2 | 32 | 1 |
| ΔΔethA2$^{CMab}$ | 16 | 1 |
| ΔΔethA2$^{CMtb}$ | 16 | 1 |
| ΔΔethA2$^{CEV}$ | 32 | 1 |

[a] Broth microdilution method was used to determine the MICs. WT: wild-type *M. abscessus*; ΔnudC: *M. abscessus nudC* knockout strain; ΔΔethA1: *ethA1* (*MAB_0985*) and *nudC* double knockout strain; ΔΔethA1$^{CMab}$, ΔΔethA1$^{CMtb}$, ΔΔethA1$^{CEV}$: ΔΔethA1 complemented with *ethA1*$^{Mab}$, *ethA*$^{Mtb}$ or with the empty vector, respectively. ΔΔethA2: *ethA2* (*MAB_013*) and *nudC* double knockout strain; ΔΔethA2$^{CMab}$, ΔΔethA2$^{CMtb}$, ΔΔethA2$^{CEV}$: ΔΔethA2 complemented with *ethA2*$^{Mab}$, *ethA*$^{Mtb}$, or with the empty vector, respectively.

**Table S2. The susceptibility of *M. abscessus* strains overexpressing *ethA1*, *ethA2*, or *ethA3* in either wild-type or *nudC* deletion backgrounds to ETH and INH.**

| ***M. abscessus* strains** | **Antibiotics/ MICs[a] (μg/mL)** | |
|---|---|---|
| | ETH | INH |
| ΔnudC$^{OEV}$ | 16 | 1 |
| ΔnudC$^{OethA1}$ | 2 | 1 |
| ΔnudC$^{OethA2}$ | 1 | 1 |
| ΔnudC$^{OethA3}$ | 2 | 1 |
| WT$^{OEV}$ | 256 | 512 |
| WT$^{OethA1}$ | 128 | 512 |
| WT$^{OethA2}$ | 256 | 512 |
| WT$^{OethA3}$ | 256 | 512 |

[a] Broth microdilution method was used to determine the MICs. ΔnudC$^{OEV}$, ΔnudC$^{OethA1}$, ΔnudC$^{OethA2}$, ΔnudC$^{OethA3}$: *M. abscessus nudC* knockout strain containing an empty vector or the vector for overexpressing *ethA1*$^{Mab}$, *ethA2*$^{Mab}$, or *ethA3*$^{Mab}$, respectively. WT$^{OEV}$, WT$^{OethA1}$, WT$^{OethA2}$, WT$^{OethA3}$: wild-type *M. abscessus* containing an empty vector or the vector for overexpressing *ethA1*$^{Mab}$, *ethA2*$^{Mab}$, or *ethA3*$^{Mab}$, respectively.

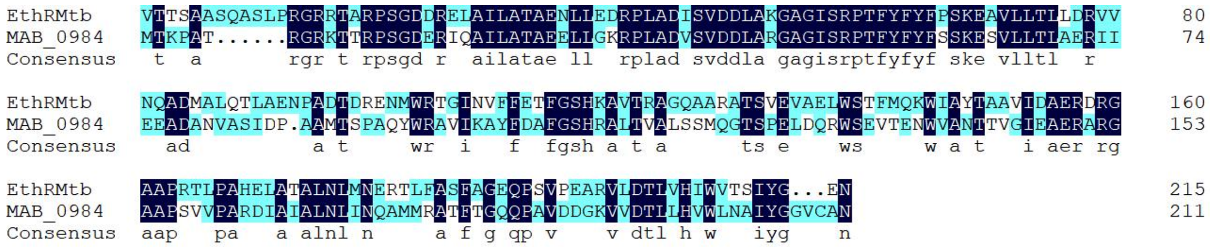


**Figure S1. Alignment of the MAB_0984 and *M. tuberculosis* EthR amino acid sequences.**

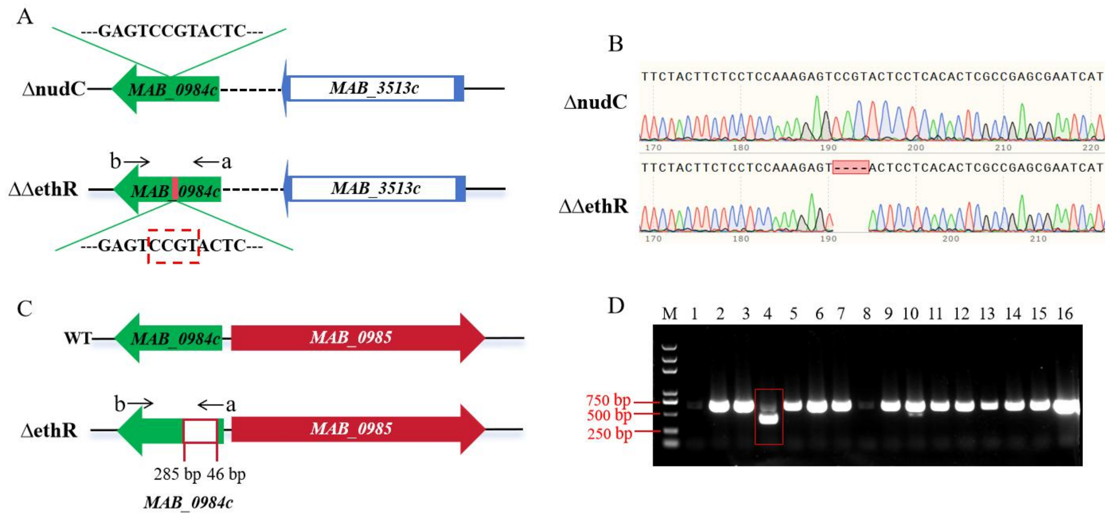


**Figure S2. Knockout of *ethR* (*MAB_0984c*) in either wild-type or *nudC* knockout strains.** (A) Schematic representation of *ethR* and *nudC* double knockout strain (ΔΔethR). A 4-bp deletion from nucleotide 191-194 was introduced, generating a frameshift mutation near the 5′ end of the gene. ΔnudC: *M. abscessus nudC* knockout strain; (B) Confirmation of the ΔΔethR knockout sequences by Sanger sequencing. (C) Schematic representation of *ethR* knockout strain (ΔethR). The ΔΔethR was constructed in the wild-type strain via an in-frame deletion of nucleotides 46-285. WT: wild-type *M. abscessus*. (D) Verification of the deletion strain by PCR and sequencing. Lane M: DNA ladder; lanes 1-15: Screening of potential ΔethR mutants by PCR using primers a and b, with lane 4 showing the product of the ΔethR; lane 16: PCR product from WT using the same primers. The expected band sizes are ~650 bp for the WT allele and ~400 bp for the deletion allele, respectively.

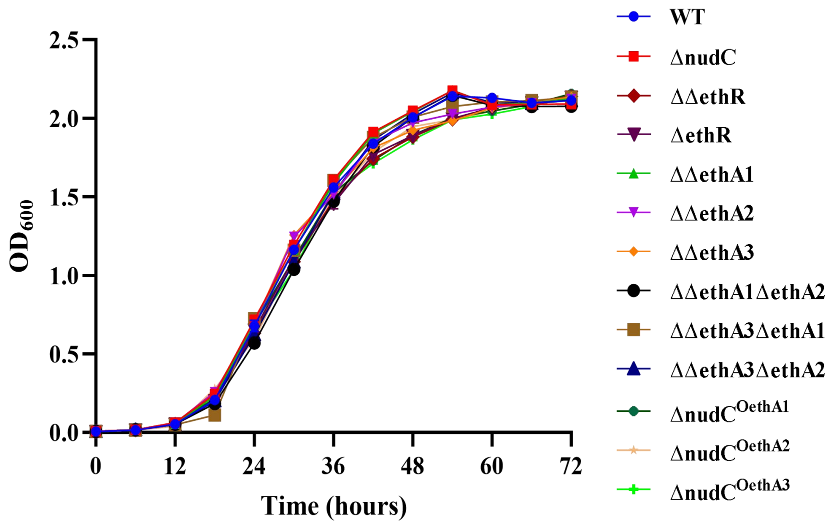


**Figure S3. Growth curves of various *M. abscessus* strains in 7H9 broth at 37°C.** WT: wild-type *M. abscessus*; ΔnudC: *M. abscessus nudC* knockout strain; ΔΔethR: *ethR* (*MAB_0984c*) and *nudC* double knockout strain; ΔΔethA1: *ethA1* (*MAB_0985*) and *nudC* double knockout strain; ΔΔethA2: *ethA2* (*MAB_0103*) and *nudC* double knockout strain; ΔΔethA3: *ethA3* (*MAB_3967*) and *nudC* double knockout strain; ΔΔethA1ΔethA2: *ethA1*, *ethA2* and *nudC* triple knockout strain; ΔΔethA3ΔethA1: *ethA3*, *ethA1* and *nudC* triple knockout strain; ΔΔethA3ΔethA2: *ethA3*, *ethA2*, and *nudC* triple knockout strain. $\Delta nudC^{OethA1}$, $\Delta nudC^{OethA2}$, $\Delta nudC^{OethA3}$: *M. abscessus nudC* knockout strain containing an empty vector or the vector for overexpressing *ethA1*$^{Mab}$, *ethA2*$^{Mab}$, or *ethA3*$^{Mab}$, respectively.

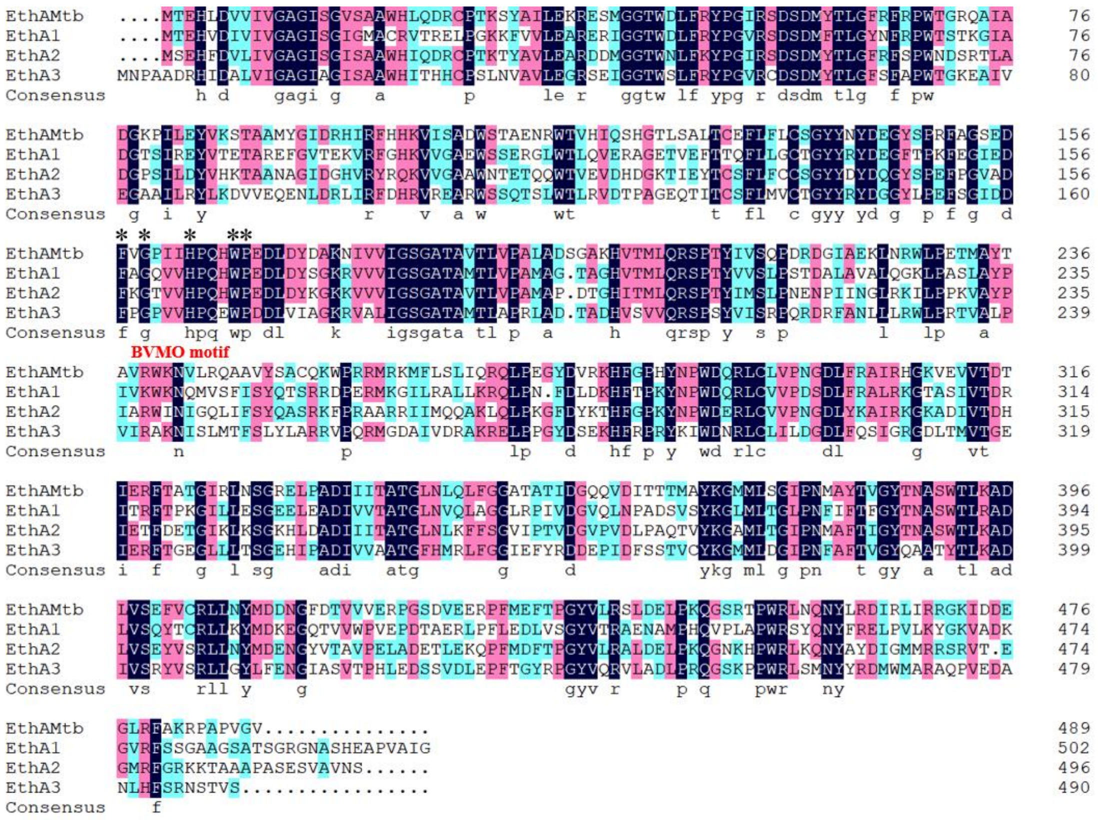


**Figure S4. Alignment of the EthA amino acid sequences from *M. abscessus* and *M. tuberculosis*.** The asterisk (*) indicates the Baeyer-Villiger monooxygenase (BVMO) motif [FXGXXXHXXXW(P/D)].

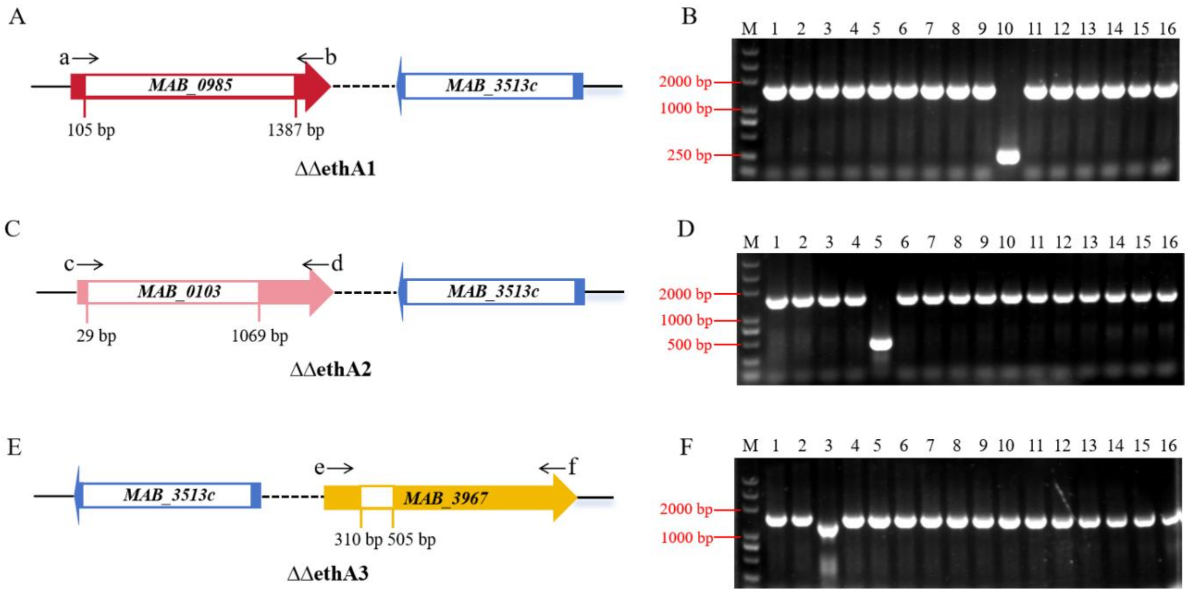


**Figure S5. Individual knockout of *ethA1* (*MAB_0985*), *ethA2* (*MAB_0103*), or *ethA3* (*MAB_3967*) in *nudC* knockout strain.** (A) Schematic representation of *ethA1* and *nudC* double knockout strain (ΔΔethA1). The ΔΔethA1 was constructed in the ΔnudC via a deletion of nucleotides 105-1387. (B) Verification of the ΔΔethA1 by PCR. Lane M: DNA ladder; lanes 1-15: Screening of potential ΔΔethA1 mutants by PCR using primers a and b, with lane 10 showing the product of the ΔethA1; lane 16: PCR product from WT using the same primers. The expected band sizes are ~1500 bp for the WT allele and ~250 bp for the deletion allele, respectively. (C) Schematic representation of *ethA2* and *nudC* double knockout strain (ΔΔethA2). The ΔΔethA2 was constructed in the ΔnudC via an in-frame deletion of nucleotides 29-1069. (D) Verification of the ΔΔethA2 by PCR. lanes 1-15: Screening of potential ΔΔethA2 mutants by PCR using primers c and d, with lane 5 showing the product of the ΔethA2; lane 16: PCR product from WT using the same primers. The expected band sizes are ~1500 bp for the WT allele and ~500 bp for the deletion allele, respectively. (E) Schematic representation of *ethA3* and *nudC* double knockout strain (ΔΔethA3). The ΔΔethA3 was constructed in the ΔnudC via a deletion of nucleotides 310-505. (F) Verification of the ΔΔethA3 by PCR. lanes 1-15: Screening of potential ΔΔethA3 mutants by PCR using primers e and f, with lane 3 showing the product of the ΔethA3; lane 16: PCR product from WT using the same primers. The expected band sizes are ~1500 bp for the WT allele and ~1200 bp for the deletion allele, respectively.

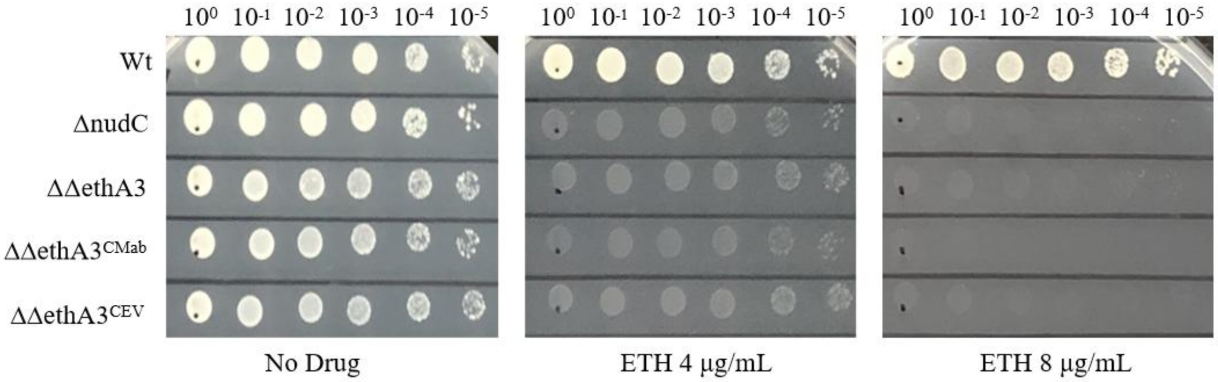


**Figure S6. Knockout of *ethA3* (*MAB_3967*) in *nudC* knockout strain increased the resistance to ETH slightly.** Drug susceptibility testing of *ethA3* and *nudC* double knockout strain to ETH. WT: wild-type *M. abscessus*; ΔnudC: *M. abscessus nudC* knockout strain; ΔΔethA3: *ethA3* and *nudC* double knockout strain; $\Delta\Delta ethA3^{CMab}$ and $\Delta\Delta ethA3^{CEV}$: ΔΔethA3 complemented with $ethA3^{Mab}$ and the empty vector, respectively.

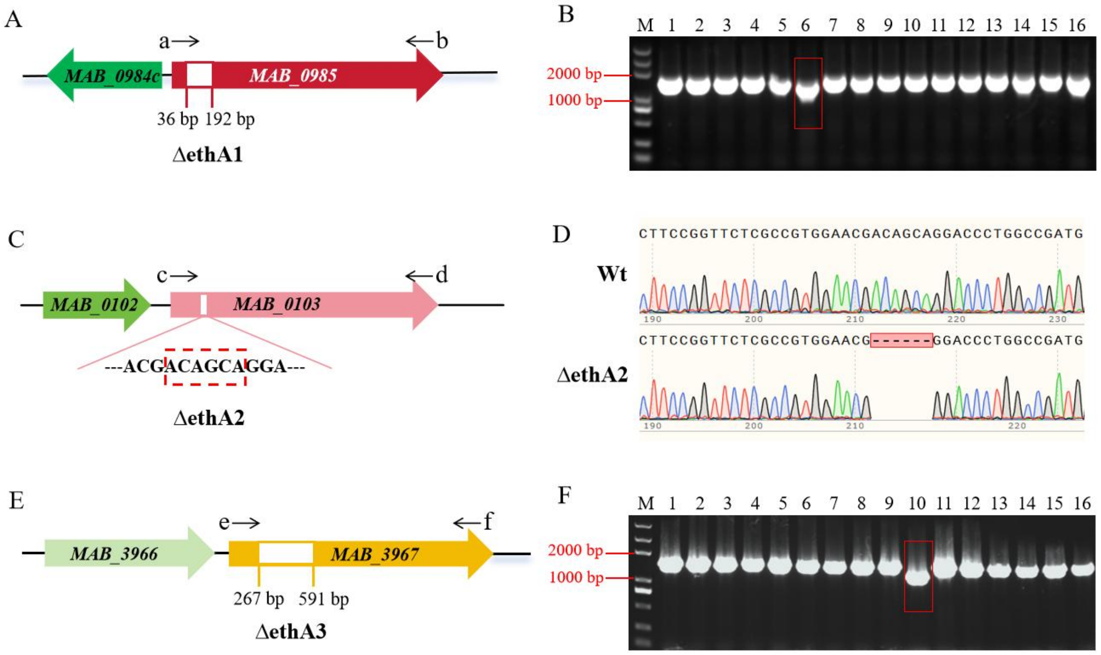


**Figure S7. Individual knockout of *ethA1* (*MAB_0985*), *ethA2* (*MAB_0103*), or *ethA3* (*MAB_3967*) in wild-type *M. abscessus*.** (A) Schematic representation of *ethA1* knockout strain (ΔethA1). The ΔethA1 was constructed in the WT via a deletion of nucleotides 36-192 in *ethA1*. (B) Verification of the ΔethA1 by PCR. Lane M: DNA ladder; lanes 1-15: Screening of potential

ΔethA1 mutants by PCR using primers a and b, with lane 6 showing the product of the ΔethA1; lane 16: PCR product from WT using the same primers. The expected band sizes are ~1500 bp for the WT allele and ~1350 bp for the deletion allele, respectively. (C) Schematic representation of *ethA2* knockout strain (ΔethA2). A 6-bp deletion was introduced to generate an in-frame deletion near the 5′ end of *ethA2*. (D) Confirmation of the ΔethA2 knockout sequence by Sanger sequencing. (E) Schematic representation of *ethA3* knockout strain (ΔethA3). The ΔethA3 was constructed in the WT via a deletion of nucleotides 267-591 in *ethA3*. (F) Verification of the ΔethA3 by PCR. lanes 1-15: Screening of potential ΔethA3 mutants by PCR using primers e and f, with lane 3 showing the product of the ΔethA3; lane 16: PCR product from WT using the same primers. The expected band sizes are ~1500 bp for the WT allele and ~1150 bp for the deletion allele, respectively.

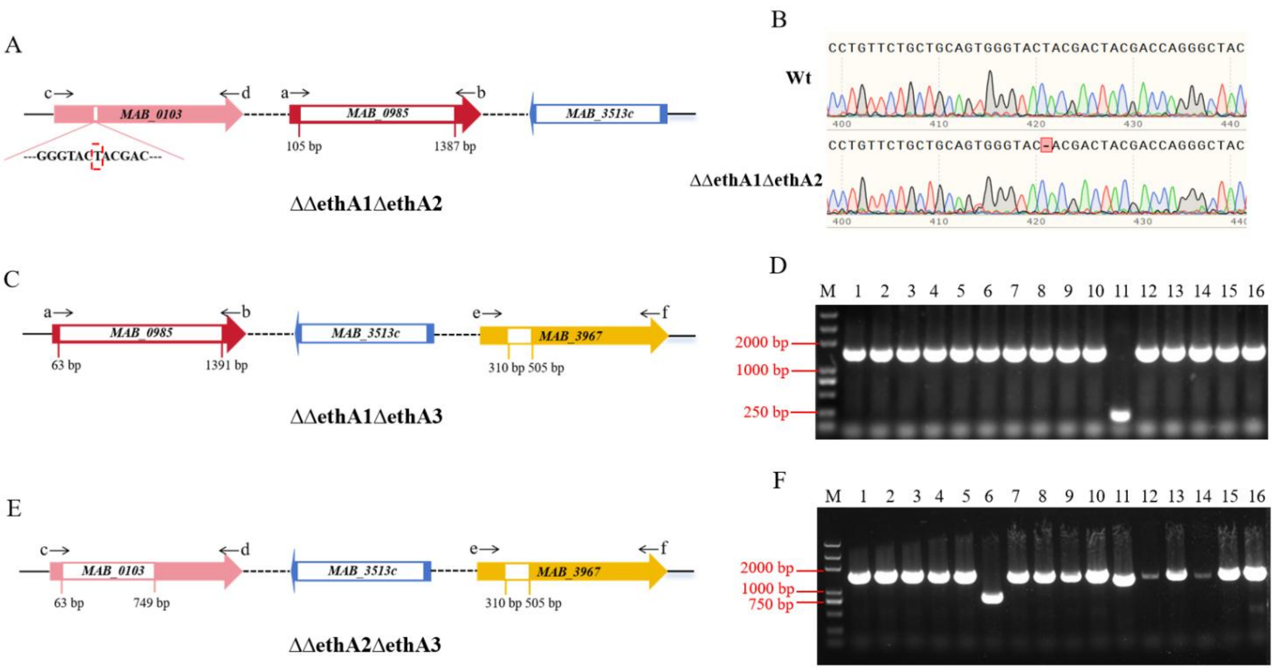


**Figure S8. Pairwise deletion of *ethA1* (*MAB_0985*), *ethA2* (*MAB_0103*), and *ethA2* (*MAB_3967*) in *nudC* knockout strain.** (A) Schematic representation of *ethA1*, *ethA2*, and *nudC*

triple knockout strain (ΔΔethA1ΔethA2). The ΔΔethA1ΔethA2 was constructed in the ΔΔethA1 via a deletion of nucleotides 421 in *ethA2*. (B) Confirmation of the ΔΔethA1ΔethA2 knockout sequence by Sanger sequencing. (C) Schematic representation of *ethA1*, *ethA3*, and *nudC* triple knockout strain (ΔΔethA1ΔethA3). The ΔΔethA1ΔethA3 was constructed in the ΔΔethA3 via an in-frame deletion of nucleotides 63-1391 in *ethA1*. (D) Verification of the ΔΔethA1ΔethA3 by PCR. lanes 1-15: Screening of potential ΔΔethA1ΔethA3 mutants by PCR using primers a and b, with lane 11 showing the product of the ΔΔethA1ΔethA3; lane 16: PCR product from WT using the same primers. The expected band sizes are ~1500 bp for the WT allele and ~200 bp for the deletion allele, respectively. (E) Schematic representation of *ethA2*, *ethA3*, and *nudC* triple knockout strain (ΔΔethA2ΔethA3). The ΔΔethA2ΔethA3 was constructed in the ΔΔethA3 via a deletion of nucleotides 63-749 in *ethA2*. (F) Verification of the ΔΔethA2ΔethA3 by PCR. lanes 1-15: Screening of potential ΔΔethA2ΔethA3 mutants by PCR using primers c and d, with lane 6 showing the product of the ΔΔethA2ΔethA3; lane 16: PCR product from WT using the same primers. The expected band sizes are ~1500 bp for the WT allele and ~800 bp for the deletion allele, respectively.

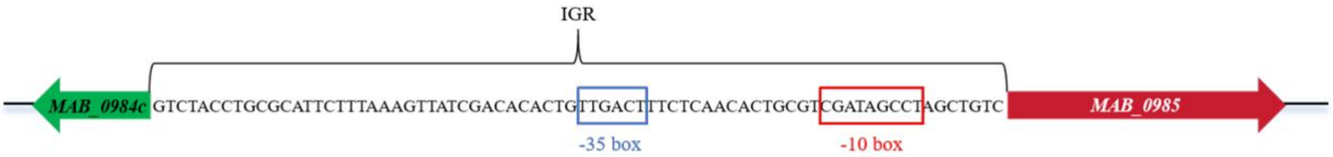


**Figure S9. Analysis of the intergenic sequence (IGR) between *ethR* (*MAB_0984c*) and *ethA1* (*MAB_0985*).**
